# *De novo* design of small cyclic peptide inhibitors from non-canonical amino acids by cofolding-guided search

**DOI:** 10.64898/2026.09.24.754019

**Authors:** Zander Harteveld, Federica Rigoldi, Chiara Toniolo, Alice Granger, Clément Gensous, Torsten Blum, Ronald Jean-Baptiste, Masa Kobayashi, Gregg Siegal, Sevan Habeshian, Lena Tagmose, Oliver Horlacher

**Affiliations:** Orbis Medicines, Copenhagen DK; ZoBio, Leiden NL

## Abstract

Cyclic peptides can engage targets beyond the reach of small molecules, but their discovery typically relies on libraries reused across targets, diversification around a known interaction motif, or computational methods limited to a handful of non-canonical building blocks. Here, we present nCycle-Forge, a simulated annealing framework that designs cyclic peptide binders by iteratively substituting building blocks and scoring each proposal by cofolding with the target. We designed, synthesized, and screened 1,472 cyclic peptides against the active-site pocket of Thrombin and the shallow p53-binding surface of MDM2 from the target sequence alone and without fixing binding motifs. The library yielded chemically diverse hits against both targets in a single round without experimental optimization, with hit rates comparable to or up to 24-fold higher than in larger experimental libraries. Purified designs reached sub- to low-micromolar potency, with the best Thrombin inhibitor at K_i_ 0.14 µM and the best MDM2 binder at EC_50_ 2.4 µM. Surface plasmon resonance confirmed direct binding, competition with orthosteric inhibitors at both sites, and weaker binding under conditions expected to linearize the designs. nCycle-Forge requires no building block-specific parameterization, conformer generation, or retraining, and provides a target-focused route to functional cyclic peptides beyond the canonical amino acid alphabet.

## Introduction

A large fraction of therapeutically relevant human proteins remains intractable to conventional drug modalities^1,2^. Targets with shallow or featureless surfaces oftentimes found on protein–protein interactions (PPIs) provide few well-defined binding pockets for traditional small molecules to bind to^3^, and even when pockets exist, achieving desired potency and selectivity across homologous proteins can be challenging^4^. Biologics such as monoclonal antibodies can engage flat epitopes with high affinity and potency but lack membrane permeability that prevents the access to the intracellular proteome and limits their therapeutic applicability^5^.

Cyclic peptides are a modality that fall in between small molecules and biologics, and thereby offer a promising solution to this long-standing gap^6^. Their ring-constrained architectures enable presentation of large, structured interaction surfaces while maintaining a compact size that can, in principle, allow for membrane permeability and oral bioavailability^6–8^. Clinically validated cyclic peptide drugs such as cyclosporine, daptomycin, zilucoplan, motixafortide, Enlicitide, Icotrokinra and various somatostatin analogues, alongside exemplars that are in late-stage clinical trials including Luna18^9,10^ illustrate the potential to combine high potency with favorable pharmacokinetics, positioning cyclic peptides as an emerging modality for targeting intracellular PPIs and other challenging targets beyond the practical reach of small molecules or antibodies.

Discovering cyclic peptide binders, however, remains challenging. Conventional approaches such as natural product mining and display technologies (e.g., phage or mRNA display) can yield potent binders but often rely on generic pre-constructed libraries that are reused across targets^11,12^. Chemically synthesized combinatorial libraries can instead be tailored to a target and provide massive chemical diversity, although even these explore only a small fraction of the accessible chemical space defined by the available amino acids (AAs) and cyclization chemistry, which can reduce the likelihood of identifying functional hits for difficult targets^13^. Moreover, advancing hits into leads requires the simultaneous optimization of potency, stability, and permeability, which remains laborious and is often difficult to achieve^14,15^.

Expanding the design space to include more diverse non-canonical amino acids (ncAAs) and linker chemistries could enable target-specific cyclic peptides with improved potency, proteolytic stability and drug-like properties^16^. ncAAs introduce chemotypes such as halogenated, fluorinated and extended heteroaromatic side chains, expanding the range and geometry of interactions available to cyclic peptides, including halogen bonding, and tailored hydrogen-bond geometries and polar–π interactions^17–19^. A suitably positioned ncAA may therefore provide binding interactions that are difficult to reproduce through canonical AAs alone. In parallel, backbone and stereochemical modifications, including D-AAs, N-methylation, α,α-disubstitution and cyclic β- or γ-AAs, can rigidify the peptide and preorganize it towards a binding-competent conformation, thereby reducing the conformational entropy penalty associated with target binding^7,16^. Hence, these expanded interaction and conformational repertoires can enable compact cyclic peptides to achieve levels of potency and selectivity more commonly associated with larger molecular modalities.

Yet, even for short cyclic peptides the accessible chemical space grows astronomically when expanding beyond the 20 canonical AAs and considering various cyclization formats (see Methods). For a linear *n*-mer peptide drawn from a building block set of size *B*, the number of possible sequences scales as *B^n^*. Accordingly, considering small cyclic peptides of length 3–8 and using *B* = 1000 ncAAs yields a total of ∼ 7.0 × 10^23^ distinct cyclic peptides, far beyond what can be accessed by experimental screening alone.

Computational approaches provide an orthogonal complement to library screening by enabling rapid *in silico* exploration of the chemical and structural space spanned by cyclic peptides. Recent advances in deep-learning structure prediction have become particularly important for this goal^20,21^. Several AlphaFold2^20^-derived methods have been adapted to cyclic peptides including approaches that encode cyclization constraints and support monomeric and complex prediction^22,23^, showing that structure prediction models can recover accurate cyclic peptide structures and peptide–target complexes. Structure prediction models have subsequently been used for design through network inversion (or “hallucination”) in which model inputs are optimized to maximize predicted confidence. This can be done by gradient-free approaches such as Monte Carlo or evolutionary search over sequences, as well as by reinforcement-learning approaches^24–28^, or by gradient-based optimization of differentiable sequence representations^29–31^. Furthermore, structure prediction methods have also been extended beyond canonical AAs by introducing pre-defined sets of ncAAs through dedicated residue representations, ncAA-containing training data or residue-specific parameterization^32,33^.

A second strategy uses diffusion-based approaches that have been used either to steer sequence-space generation with structure-prediction confidence during denoising^34^ or to generate cyclic candidate backbones for subsequent sequence design by inverse folding^35–38^. In RFpeptides^35^ this has produced experimentally validated macrocyclic binders. In parallel, approaches built on physics-based scoring functions have yielded *de novo* peptide macrocycles, including membrane-permeable examples^39–45^. Collectively, these advances show that structure-based computational design can generate accurate and, in some cases, bioactive and functional cyclic peptides.

The latest implementations of all-atom structure prediction models represent small molecules a the atom-level, enabling the cofolding of protein–ligand complexes directly from chemical structure and protein AA sequence^46,47^. These cofolding models have subsequently been adapted to the prediction and design of cyclic peptides containing ncAA^48–51^.

Despite this progress, many cyclic peptide design methods either operate over the 20 canonical AAs, use relatively small predefined ncAA alphabets, or support a restricted set of cyclization chemistries. Other approaches make use of computationally generated conformer ensembles, whose generation can itself be costly^52^.

A complementary challenge is therefore to search large, chemically explicit building block collections while retaining the ability to vary building blocks and linker chemistry without model-specific residue parameterization or retraining.

Molecular graph-based representations used by all-atom cofolding models could enable a route to predict ncAA-containing cyclic peptides by inputting them as graphs and thereby avoiding the need for special residue representation or parametrization, or the need to retrain or fine-tune the model for each newly introduced building block.

Many ncAAs incorporate chemotypes that are common in small molecule ligands—the average ligand in the Protein Data Bank (PDB)^53^ has roughly 50% AA character^54^—and the geometries of protein–small molecule contacts can often be approximated by protein–AA contacts^55,56^. Importantly, AlphaFold2, despite being trained solely on monomeric protein structures, can accurately predict protein–peptide complexes without peptide co-evolutionary information or examples of peptide complexes in its training set^57–59^.

These observations suggest that structure-prediction models can encode transferable information about local interaction geometries beyond what they’ve seen during training. We therefore reasoned that cofolding models trained on the PDB, including protein– small molecule complexes, may generalize to a certain extent to ncAA and capture useful interaction patterns and conformational preferences.

Motivated by this hypothesis, we represent cyclic peptides as atom-level graphs rather than as AA sequences. Because inference is fast for modestly sized proteins, we embed a cofolding model (here Boltz^60,61^) within a gradient-free search algorithm, simulated annealing, that iteratively proposes ncAA modifications and selects candidates that improve the model’s predicted confidence scores and thereby explores the chemical space toward high-scoring designs. The optimization procedure is flexible, readily allowing additional design objectives to be incorporated by augmenting the objective score with user-defined terms (e.g., penalties for Rule-of-5^62^ violations), enabling co-optimization of multiple parameters.

Our method, dubbed nCycle-Forge, provides a fast, programmable framework for the *de novo* design of small cyclic peptides composed of ncAAs from the target protein sequence alone.

## Results

### nCycle-Forge enables tunable design of cyclic peptide binders

We first asked whether Boltz2 could recover the receptor-bound poses of cyclic peptides input as 2D chemical graphs. On a curated set of 240 experimentally solved macrocycle–target complexes^63^ with pocket constraints applied so that the task was restricted to local pose recovery (Sup. Fig. 1; Sup. Methods), Boltz2 placed 85% of macrocycles (≤85 atoms) below 2 Å RMSD and 26% below 0.5 Å, with failures dominated by flipped or mispositioned poses (Sup. Fig. 1E,F). However, relatively few experimentally determined small cyclic peptide–target complexes were deposited after the model’s training cutoff, limiting the extent to which this retrospective analysis can establish generalization to unseen molecules. Using the model generatively provides a more direct test, since designed molecules can be assessed both for chemical novelty and for function.

We therefore set out to determine whether cofolding could serve as an engine for the *de novo* design of cyclic peptides against specific protein targets. We formulated design as an iterative search over a surrogate ligand–target interaction landscape defined by Boltz2 output scores (Fig. 1A; Methods). Starting from randomly initialized cyclic peptides with a specified length and cyclization format, we repeatedly introduced single building block substitutions (including ncAAs and linkers) drawn from a predefined set, cofolded each proposal with a target protein sequence and selected mutations using a simulated annealing schedule (Fig. 1B,C; Methods). Early in the trajectory, occasional acceptance of lower-scoring proposals promotes exploration of the sequence space. As temperature decreases, the procedure becomes progressively greedier, and thereby effectively approaches hill-climbing on the surrogate landscape (Methods).

**Figure 1.**
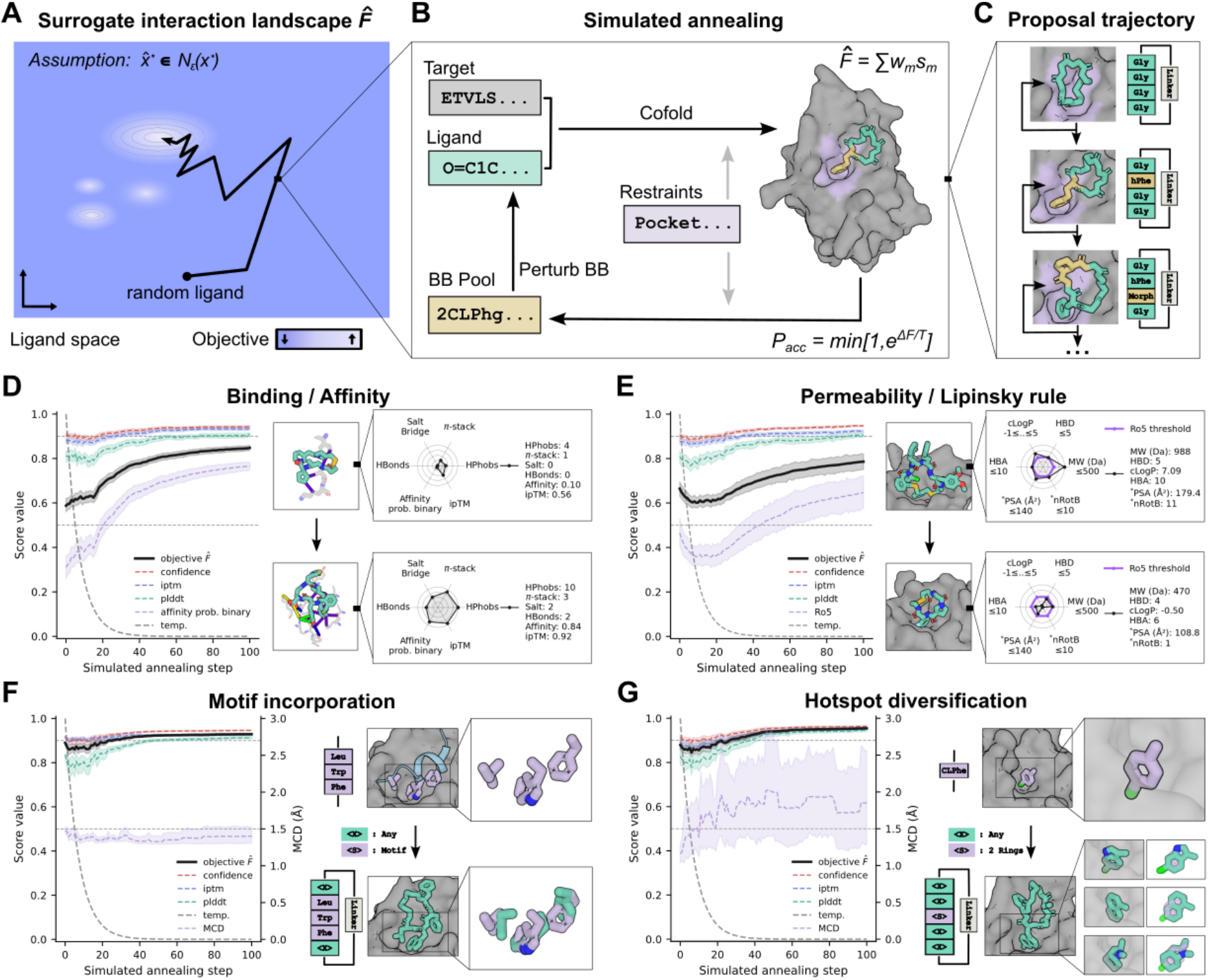
Overview of nCycle-Forge. **A:** Conceptual surrogate fitness landscape *F̂* used for ligand optimization. Although *F̂* may differ from the true interaction landscape *F*, ascent on *F̂* is assumed to enrich candidates near the true interaction peak (see Methods). A representative stochastic trajectory for a ligand is shown. **B:** Workflow of nCycle-Forge. At each step, the cyclic peptide is locally perturbed by a single building block substitution, cofolded with the target, and scored using a weighted surrogate objective. Proposed mutations are accepted using a Metropolis criterion under an exponentially decaying temperature schedule. **C:** Example optimization trajectory showing iterative building block substitutions and corresponding cofolded poses, illustrating progressive improvement in predicted binding geometry and surrogate score. **D:** Mean objective score improvement over 100 independent optimization trajectories using an objective combining confidence, ipTM, ligand pLDDT and binary probability affinity prediction, plotted across annealing steps. **E:** Incorporation of beyond Rule-of-5 permeability constraints into the objective. Trajectories simultaneously improve the objective score while reducing violations of beyond Rule-of-5 related thresholds. **F:** Motif integration during optimization. Native binding interaction motifs are enforced via distance restraints, enabling motifs to be scaffolded into cyclic peptide ligands while improving the predicted score. The mean center deviation (MCD) is defined as the Euclidean distance between the geometric centers (centroids) of the reference motif ligand and the designed hotspot residue after superimposing receptor structures using their Cα atoms. MCD was not included in the objective score and was used only for monitoring during the optimization trajectories. **G:** Hotspot diversification. Shape-matched but chemically distinct building blocks are explored at defined interaction hotspots, sampling alternative substituents while preserving the predicted binding mode.

Using a simple objective that uniformly combines confidence score, ipTM, mean ligand pLDDT and the predicted (binary probability)-affinity module output (Fig. 1D and Methods), nCycle-Forge rapidly generated high-scoring cyclic peptides within short runs (Fig. 1D). Across 100 independent trajectories, the objective improved consistently over the course of the optimization, with high-scoring candidates typically emerging within as few as 100 steps (Fig. 1D). We observed a median improvement of ∼0.26 in objective score (from 0.57 to 0.84) and ∼0.45 in predicted affinity (from 0.30 to 0.75) with nearly all designs surpassing the 0.5 threshold (Sup. Fig. 2A). The fraction of trajectories exceeding a high objective score threshold of 0.8 increased from 6% at initialization to 80% after optimization, indicating an efficient search despite the large discrete design space (Sup. Fig. 2A).

A key advantage of the simulated annealing formulation is that it supports tunable design by directly augmenting the objective with user-defined penalties and constraints (Methods). For example, to bias designs toward more drug-like physicochemical properties, we added a penalty term capturing violations of the Rule-of-5 (Ro5) descriptors (Methods). This shifted the optimization trajectory toward designs with improved Ro5 profiles while maintaining high predicted confidence and objective score (Fig. 1E, Sup. Fig. 2B), demonstrating that penalties can be readily integrated and co-optimized with the confidence metrics.

We then asked if we could extend our method to scaffold known interaction motifs i.e., embed them into a cyclic peptide context while maintaining their 3D geometry intact. Starting from residue motifs derived from the native MDM2 ligand, the p53 transactivation domain (PDB 4HFZ; motif residues F19, W23, L26), we derived sets of distances from the motifs to the target structure and used these as steering constraints during each cofolding step over the trajectory (Methods). Under these constraints, nCycle-Forge generated cyclic peptides that preserved the intended hotspot geometry while building a cyclic scaffold around it, thus effectively “cyclizing” motifs into compact cyclic peptide ligands (Fig. 1F).

Finally, we asked whether the same framework could support hotspot diversification. Rather than simply preserving a motif, we implemented a 3D shape-matching procedure to identify compatible substitutions that maintain the overall geometry and shape while altering the functional group. We applied this diversification to a known MDM2 binding motif (PDB 1T4E; benzodiazepinedione inhibitor, with the central chlorophenyl as motif) by scanning a set of ncAAs that resemble the native moieties. We found that nCycle-Forge could identify alternative ncAAs that fit the native geometry and have similar shapes (Fig. 1G). This enabled exploration of alternative pharmacophores while preserving key contacts and introducing new potential interactions at the interface (Fig. 1G).

Together, these examples demonstrate the versatility of nCycle-Forge across distinct cyclic peptide design tasks. By decoupling programmable, constraint-aware optimization from the underlying cofolding engine, nCycle-Forge can, in principle, be paired with any structure-prediction method that outputs a complex structure and associated confidence or scoring metrics, providing a general steering framework for structure-based cyclic peptide design.

### Computational design, synthesis and screening of 1,472 cyclic peptides against Thrombin and MDM2

Encouraged by the computational results, we sought to experimentally test whether the nCycle-Forge generated cyclic peptides would be truly functional—that is, whether they bind and inhibit their intended targets—or if they instead are high-scoring but non-functional *in silico* artifacts.

To this end, we designed a library of cyclic peptides targeting either Thrombin or MDM2. Thrombin is a trypsin-like serine protease that plays a central role in blood coagulation and has a deep, well-defined catalytic active-site pocket that is readily assayed by substrate turnover (Fig. 2A). MDM2 is an E3 ubiquitin ligase and negative regulator of the tumor suppressor p53. Inhibiting MDM2 can restore p53 activity in tumors that retain wildtype p53. Structurally, MDM2 recognizes an amphipathic α-helical segment of the p53 transactivation domain, dominated by the hydrophobic hotspot residues Phe19, Trp23, and Leu26, which insert into a comparatively broad hydrophobic cleft on MDM2 (Fig. 2C).

**Figure 2.**
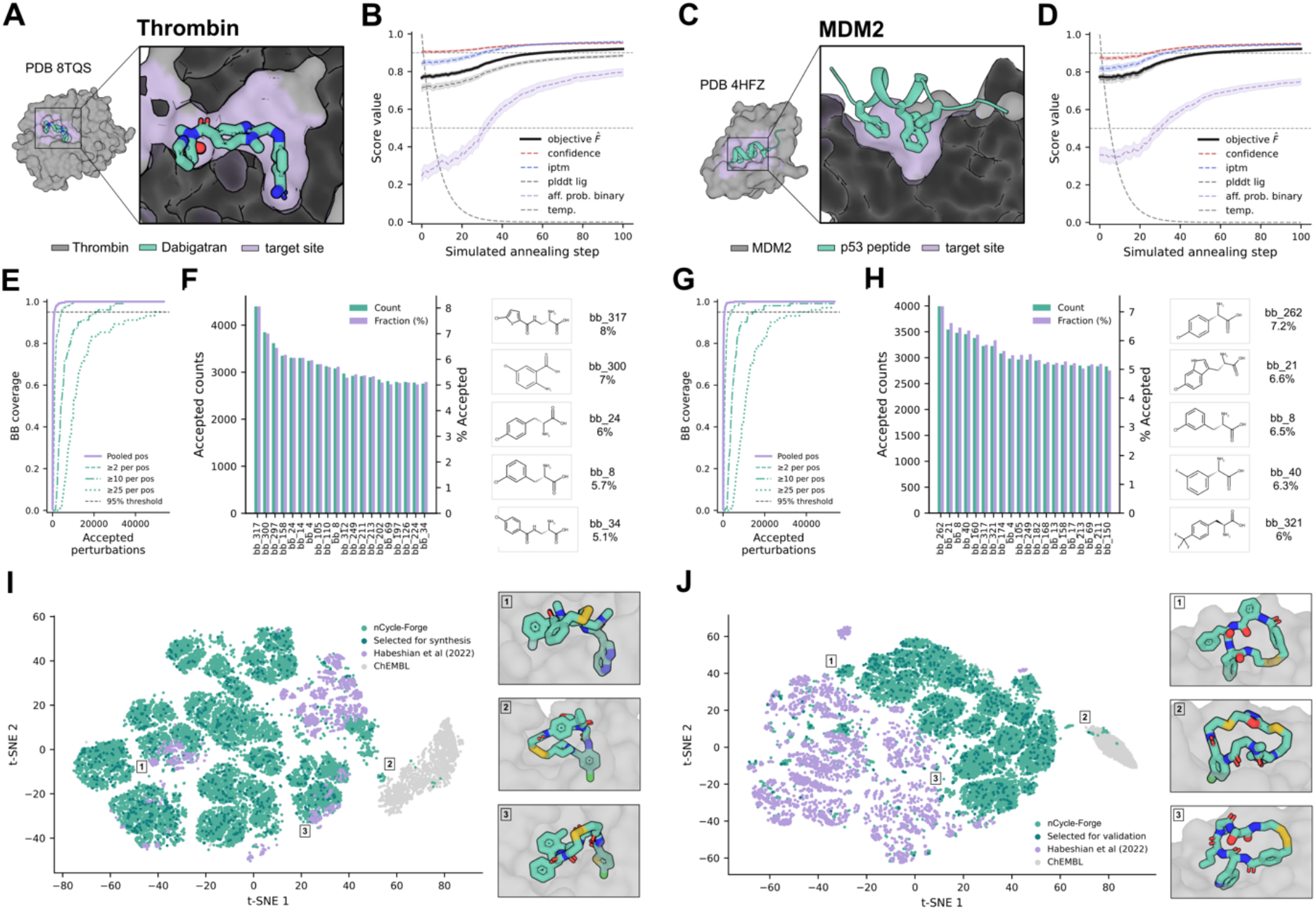
Computational cyclic peptide library targeting Thrombin and MDM2. **A:** Structure of Thrombin (gray surface) in complex with the inhibitor Dabigatran (PDB 8TQS). **B:** nCycle-Forge optimization trajectory for Thrombin, averaged over ∼600 independent trajectories of 100 steps each. **C:** Structure of MDM2 (gray surface) in complex with its native binding partner p53 (PDB 1YCR). **D:** nCycle-Forge optimization trajectory for MDM2, averaged over ∼600 independent trajectories of 100 steps each. **E,G:** Building block coverage aggregated across trajectories, showing that each building block was accepted nearly 25 times per position for each target, indicating thorough sampling at the building block level. **F,H:** The 20 most frequently accepted building blocks, highlighting ncAAs enriched in designs for Thrombin or MDM2. **I,J:** Chemical diversity of designed cyclic peptides visualized by t-SNE embedding of MHFP fingerprint distances (Methods). Designs span a broad, different region of chemical space compared to prior macrocyclic libraries and ChEMBL small molecules.

The two targets pose contrasting design challenges. Thrombin presents a geometrically restrictive enzymatic pocket, whereas MDM2 presents a broader, more open PPI groove that requires productive placement of several anchor groups over a larger interaction surface (Fig. 2A,C).

To explore how different model and objective choices affect design, we ran nCycle-Forge in three different configurations: (i) Boltz1 (v1.0.0) and (ii) Boltz2 (v2.1.1), both using a uniformly weighted objective combining confidence, ligand pLDDT and ipTM; and (iii) Boltz2+affinity, which additionally incorporates the binary probability affinity module as part of the objective.

We initialized the searches with head-to-tail cyclic peptides spanning 3-6 ncAAs and one of 6 linkers. Because our synthesizer supports up to ∼102 building blocks whereas our collection contains >1,000, we used a two-stage approach. First, we performed an exploratory design round using the full ncAA set to identify building blocks enriched among high-scoring solutions (Sup. Fig. 3A,C–F). Second, we carried out production runs restricted to the 102 enriched ncAAs from the exploration rounds (Sup. Fig. 3B).

Across the production simulations for Thrombin and MDM2, we performed ∼600 independent trajectories at 100 steps each per target, totaling to ∼120,000 cofolding evaluations (61,098 for Thrombin and 61,200 for MDM2) (Fig. 2B,D). Importantly, each building block was sampled nearly 25 times per position for each target, indicating broad sampling at the building block level (Fig. 2E,G). Strikingly, a subset of building blocks was repeatedly enriched among top-scoring designs, and enrichment patterns differed between Thrombin and MDM2 despite using the same 102 ncAAs, which is consistent with target-specific design solutions (Fig. 2F,H).

We next assessed the structural and chemical diversity of the designed cyclic peptides. Compared to a previously reported systematic macrocycle library^13^ and with small molecules sourced from ChEMBL^64^, nCycle-Forge designs occupied a different region of the embedded chemical space for both targets, indicating that the method samples a diverse set of cyclic peptides rather than converging to a narrow motif family (Fig. 2I,J).

Because the automated synthesizer accommodates four 384-well plates, we down-sampled to a synthesis-ready set of 1,472 designs split evenly between Thrombin and MDM2 designs (736 per target; leaving 64 empty wells as buffers).

Candidates were selected by first applying a Rosetta ddG filter (ddG < 0) (Methods), removing designs with an unfavorable change in free energy when complexed. The remaining designs were ranked by the nCycle-Forge objective score while enforcing diversity through a Jaccard distance criterion between the sequences (Jaccard distance > 0.9) to avoid over-representing near-duplicates (Methods).

Initial synthesis tests revealed a subset of 13 ncAAs associated with systematic coupling or solubility failures, reducing product purity (Sup. Fig. 4A). To improve the quality of the final library, we capped the frequency with which these problematic building blocks were used.

We synthesized the library in a 384-well format by solid-phase synthesis (Fig. 3A; Methods). LC–MS analysis of 88 selected crude library products indicated a median purity of ∼76% (Fig. 3B,C; Sup. Fig. 4B; Methods). We considered this purity sufficient for direct screening without chromatographic purification and therefore tested the crude library for inhibition of Thrombin activity and disruption of the MDM2:p53 interaction.

**Figure 3.**
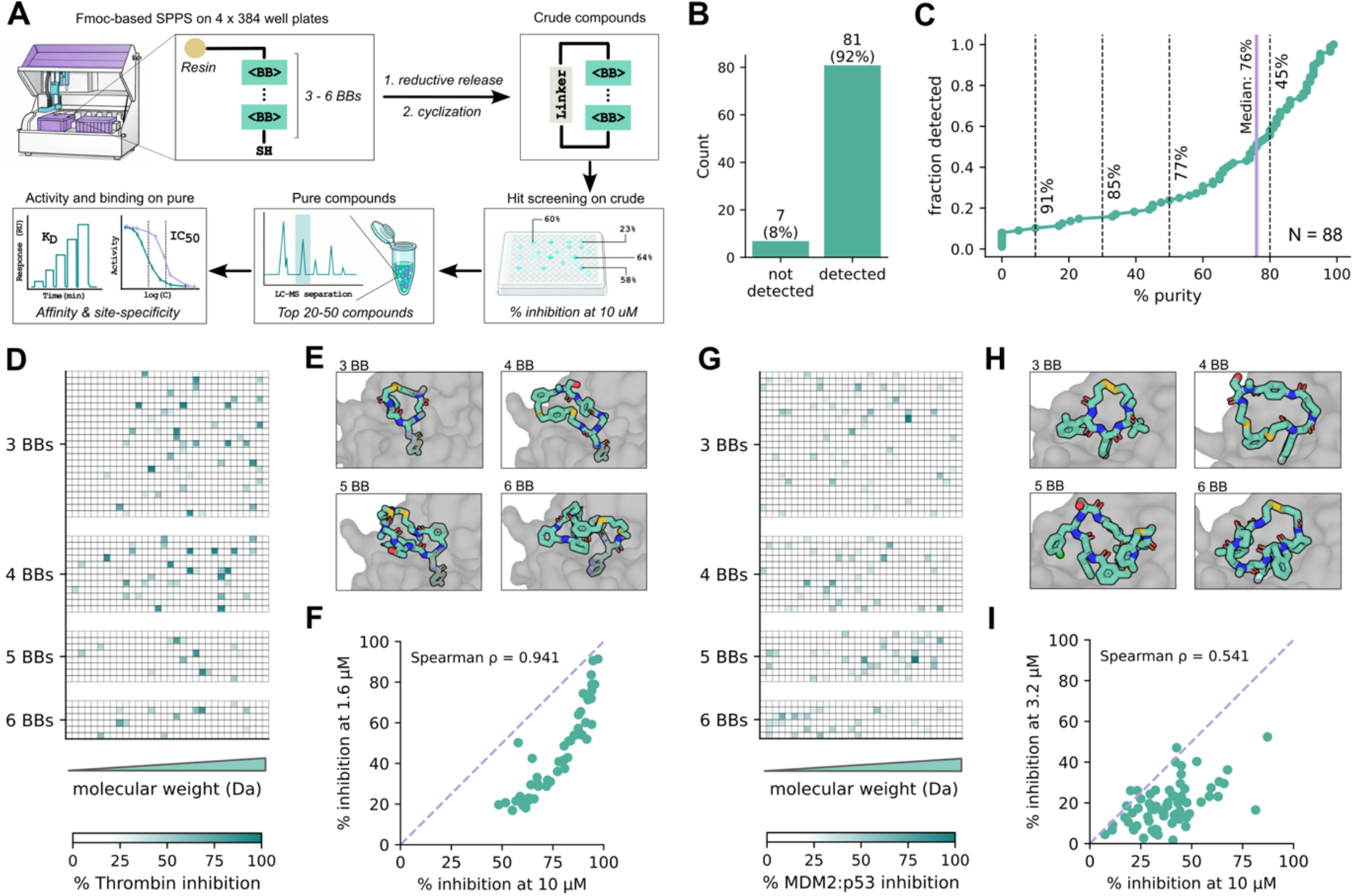
Synthesis and screening of the crude library. **A:** Experimental workflow for library synthesis and screening. Linear peptides were synthesized by solid state peptide synthesis in 384-well plates and cleaved by reductive release, followed by cyclization. Primary hit identification was performed directly on crude material, followed by hit confirmation through re-synthesis, purification, and orthogonal validation by biochemical assays and SPR. **B,C:** Purity of a selected subset of 88 crude cyclic peptides (Sup. Fig. 4 and Methods). Of these, 81 compounds were detected at the expected mass and median crude purity was 76%. **D:** Heatmap showing inhibition of Thrombin enzymatic activity for each compound in the crude library. Results are stratified by cyclic peptide length. **E:** Most active crude Thrombin hits by cyclic peptide length. **F:** Re-screening of crude Thrombin hits (>50% inhibition) at two concentrations to assess concentration dependence. **G:** Heatmap of the inhibition of MDM2-p53 interaction in HTRF by each compound of the crude library. Results are stratified by length of the cyclic peptides. **H:** Most active crude MDM2 hits stratified by cyclic peptide length. **I:** Re-screening of crude MDM2 hits (>25% inhibition) at two concentrations to assess concentration dependence.

For Thrombin, we used a fluorogenic substrate turnover assay (Methods). Screening the crude designs at 10 µM identified inhibitors spanning a range of potencies, with hits across all cyclic peptide lengths tested (Fig. 3D,E). 35 out of 736 designs (∼5%) inhibited Thrombin by more than 50%. Interestingly, undirected screening of comparably prepared crude cyclic peptide libraries against Thrombin, that is libraries not computationally designed or enriched for Thrombin binding, at the same screening concentration and inhibition threshold (10 µM, >50% inhibition) previously yielded hits in only 0.2% (9 of 4,608) and 0.9% (73 of 8,448) of wells^13,14^, corresponding to 24- and 6-fold lower hit rates. To assess reproducibility and concentration dependence and prioritize hits, the most active crude hits were re-tested at 10 and 1.6 µM (Fig. 3F). 34 of the 35 Thrombin designs retained >50% inhibition at 10 µM, and inhibition measurements were strongly rank-correlated across the two concentrations (Spearman ρ = 0.96), consistent with reproducible activity and suggesting that several candidates may represent genuinely potent Thrombin inhibitors.

For MDM2, an HTRF competition assay reporting disruption between MDM2 and p53 was used to identify competitive binders (Methods). Across the crude library we observed a broad distribution of inhibition values, with a larger fraction of weak inhibitors and few strong candidates (compared to Thrombin) (Fig. 3G,H). 88 of the 736 MDM2 designs (12%) reached >25% inhibition in the primary crude screen. Because single-point screening of crude material is noisy, the 40 most active MDM2 designs were re-tested at 10 and 3.2 µM (Fig. 3I). 34 retained >25% inhibition at 10 µM, corresponding to a confirmed hit rate of 4.6% (34 of 736), and the rank correlation between concentrations was moderate (Spearman ρ = 0.57), with a subset of compounds retaining inhibition and showing concentration dependence. A previous target-tailored combinatorial cyclic peptide library, in which every scaffold contained a p53-mimicking tryptophan or phenylalanine hotspot, yielded a comparable hit rate against the same interaction^13^ (4.0% at >25% probe displacement after excluding one building block that displaced probe across all scaffolds and was flagged in that study as assay-interfering). nCycle-Forge matched this hit rate after confirmation and without fixing a hotspot.

Computational design of a chemically diverse, target-focused library, coupled with automated synthesis and direct screening of crude products enabled rapid identification of diverse cyclic peptide hits against both an enzymatic pocket and a PPI surface highlighting the practicality and versatility of nCycle-Forge for hit discovery across distinct binding-site geometries.

### Designed cyclic peptides show potent Thrombin inhibition

From the crude Thrombin screen, 23 compounds with promising inhibitory activity were selected for tube-scale re-synthesis and purification (Methods; Sup. Table 1; Sup. Fig. 5.1–5.3D; Sup. Fig. 5.4E). We quantified inhibitory potency for purified fractions using the fluorogenic substrate turnover assay across a 16-point concentration series and three independent replicates (Methods). Of the 27 purified fractions tested, 24 retained concentration-dependent inhibitory activity, with a median K_i_ of 2.7 μM (Fig. 4D; Sup. Fig. 5.1–5.3D; Sup. Fig. 5.4E; Sup. Table 1). Across fractions of ≥80% LC–MS purity, K_i_ values spanned 0.14–12.9 μM, with thbn_cp11 the most potent (K_i_ = 0.14 μM, 90% purity). Two lower-purity fractions, thbn_cp19a (K_i_ = 0.13 μM, 52% purity) and thbn_cp5 (K_i_ = 0.23 μM, 77% purity), showed comparable potency but were not carried into biophysical characterization. Although the designed inhibitors did not reach the low-nanomolar potency of Dabigatran (K_i_ = 1.1 nM) (Sup. Table 1), the most active fractions were low- to submicromolar inhibitors, with several of them within an order of magnitude of the previously reported macrocyclic Thrombin inhibitor M1^13^ (K_i_ = 28 nM) (Sup. Table 1; Sup. Fig. 5.4F), which was identified by synthesizing and screening a combinatorial library of roughly 20,000 macrocycles. Activity measured after purification was broadly consistent with the initial crude-screen prioritization, supporting direct screening of crude products as a useful strategy for identifying and triaging active cyclic peptide inhibitors.

**Figure 4.**
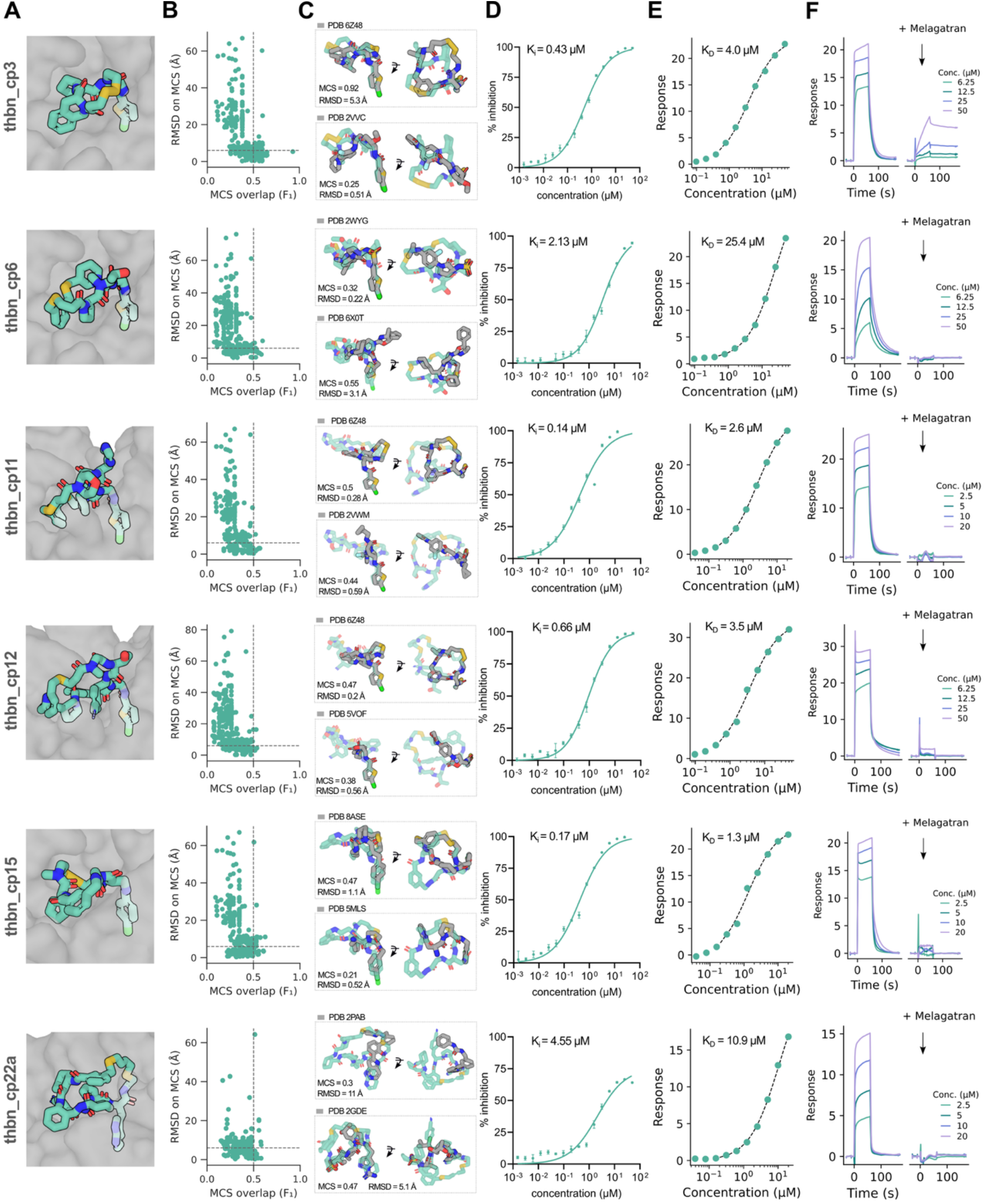
Computational and biochemical data for top *de novo* Thrombin inhibitors. **A:** Structural models of designed cyclic peptide designs (colored and stick representation) bound to the Thrombin active site pocket (shown as gray surface). **B:** Structural similarity analysis comparing each cyclic peptide design to ligands observed in Thrombin-bound structures from the PDB. Foldseek was used to identify relevant Thrombin-ligand complexes from the PDB. Retrieved structures were aligned through the Thrombin protein, followed by MCS identification between each design and PDB ligand. The RMSD of the shared MCS fragment was then calculated. Scatter plots show MCS overlap fraction (reported as the F1 score) on the x-axis and MCS-fragment RMSD on the y-axis. Points with high MCS overlap and low RMSD represent ligands with the closest structural and spatial similarity to the designed cyclic peptides. **C:** Examples of PDB ligands with high MCS overlap or low RMSD to the designs. **D:** Dose-response curves from Thrombin inhibition assay, showing concentration-dependent inhibition. **E:** Equilibrium SPR binding analysis showing concentration-dependent binding responses and corresponding fitted dissociation constants (K_D_) for interaction with Thrombin. **F:** Competitive SPR sensorgrams in the presence and absence of Melagatran. Reduced peptide binding in the presence of Melagatran indicates competition for the Thrombin active site and supports binding of the designs to the same or an overlapping pocket as Melagatran.

We next examined whether selected active designs recapitulated interaction motifs observed in experimentally determined Thrombin–ligand complexes. Structural models placed representative cyclic peptides in the Thrombin active-site pocket, with several designs positioning hydrophobic aromatic or heteroaromatic ncAA side chains deep within the substrate-recognition pocket (Fig. 4A). For the top designs, we structurally aligned related PDB complexes via the Thrombin protein and calculated the maximum-common-substructure (MCS) overlap and root-mean-square-deviation (RMSD) between PDB ligands and either the designed cyclic peptides or individual building blocks (Fig. 4B; Methods).

For design thbn_cp3, this analysis identified a chemically similar crystallographic cyclic peptide ligand (PDB 6Z48), with an MCS overlap of 92% and an MCS RMSD of approximately 5 Å (Fig. 4C; Sup. Fig. 7). Inspection of the aligned structures showed that both ligands present a chlorothiophene moiety that docks into the Thrombin pocket in a similar orientation, while the overall macrocyclic scaffolds are comparable in size but differ in their building block composition (Fig. 4C; Sup. Fig. 7). For other designs containing chlorothiophene motifs, including thbn_cp3, thbn_cp6, thbn_cp11 and thbn_cp12, the same analysis identified related substructure matches to small molecule Thrombin ligands, indicating that this chemotype is a recurrent motif among known Thrombin inhibitors (Fig. 4A,B,C; Sup. Fig. 7–10). Additional designs positioned chlorophenyl moieties (e.g., thbn_cp15) into the pocket, another motif observed among Thrombin-bound small molecules (Fig. 4A,B,C; Sup. Fig. 11).

In addition to these known motifs, a selected design thbn_cp22a positioned an azatryptophan motif into the Thrombin pocket, for which we did not identify an equivalent motif among the analyzed PDB ligands (Fig. 4A,B,C last row; Sup. Fig. 12). This moiety was not represented among the retrieved Thrombin-bound ligands, indicating that nCycle-Forge also sampled outside the recurrent chemotypes (Sup. Fig. 12). Overall, the observed similarities were primarily local rather than whole-molecule matches, indicating that selected designs incorporated known pocket-engaging chemotypes into larger and chemically distinct cyclic peptide scaffolds while also sampling alternative interaction motifs.

To quantify direct binding, we analyzed a subset of 17 purified fractions by surface plasmon resonance (SPR). Human α-Thrombin was immobilized, and cyclic peptides were injected as concentration series with top concentrations adjusted according to NMR-derived solubility limits (Methods; Sup. Table 1). Sensorgrams were fitted using standard kinetic or steady-state models to estimate binding affinities, with Melagatran and M1 included as positive controls (Methods). The estimated K_D_ values ranged from 1.3 to 84.3 μM, with a median K_D_ of 12.6 μM (Fig. 4E; Sup. Fig. 5.1–5.3E,F; Sup. Fig. 5.4F,G, Sup. Table 1). For six compounds, full kinetic fitting was feasible and revealed relatively fast on- and off-rates (Sup. Fig. 5.1–5.3E; Sup. Fig. 5.4F, Sup. Table 1). All tested compounds produced reproducible binding signals above reference, consistent with direct binding to Thrombin. SPR-derived binding affinities correlated with the corresponding K_i_ values from the enzymatic inhibition assay (Spearman ρ = 0.74), indicating that stronger direct binding translated into increased inhibitory potency (Sup. Table 1).

To confirm that the compounds bind the intended Thrombin active-site pocket, we performed competitive SPR experiments using Melagatran, a well-characterized orthosteric Thrombin inhibitor. Peptide binding was measured at multiple concentrations in the absence or presence of saturating Melagatran. For the tested compounds, SPR responses were strongly reduced or abolished in the presence of Melagatran, indicating that peptide binding is competitive and therefore occurs at the same or an overlapping active-site pocket (Fig. 4F; Sup. Fig. 5.1–5.3G; Sup. Fig. 5.4H).

The biochemical inhibition data, predicted active-site poses for representative designs, ligand-fragment comparisons for selected examples, direct SPR binding, and Melagatran competition data support that the designed cyclic peptides engage Thrombin directly and likely occupy the intended active-site pocket.

### Designed cyclic peptides disrupt the MDM2:p53 interaction

A total of 18 compounds that showed promising inhibition in the crude MDM2 screen were selected for tube-scale re-synthesis and purification (Methods; Sup. Table 2; Sup. Fig. 6.1–6.3A–D). In total, 23 purified fractions were isolated, reflecting chromatographic separation of multiple fractions for a subset of compounds, likely corresponding to separable stereoisomeric or closely related product species (Sup. Table 2). All purified fractions were characterized by analytical HPLC/UPLC and LC–MS prior to biochemical and biophysical testing (Methods; Sup. Fig. 6.1–6.3B–D).

We next quantified disruption of the MDM2:p53 interaction using a 16-point homogeneous time-resolved fluorescence (HTRF) competition assay (Methods). Among the active purified fractions, apparent EC_50_ values ranged from 2.4 to 50.3 μM (Fig. 5D; Sup. Fig. 6.1 –6.3E; Sup. Table 2). The most potent designs came within roughly an order of magnitude of the small molecule inhibitor Nutlin-3A which showed an EC_50_ of 0.22 μM under the same assay conditions. Several designs were more potent than the previously reported cyclic peptide M6^13^ which showed an EC_50_ of approximately 32 μM (Sup. Table 2; Sup. Fig. 6.3E). These results confirmed that the designed cyclic peptides can functionally disrupt the MDM2:p53 interaction after purification.

**Figure 5.**
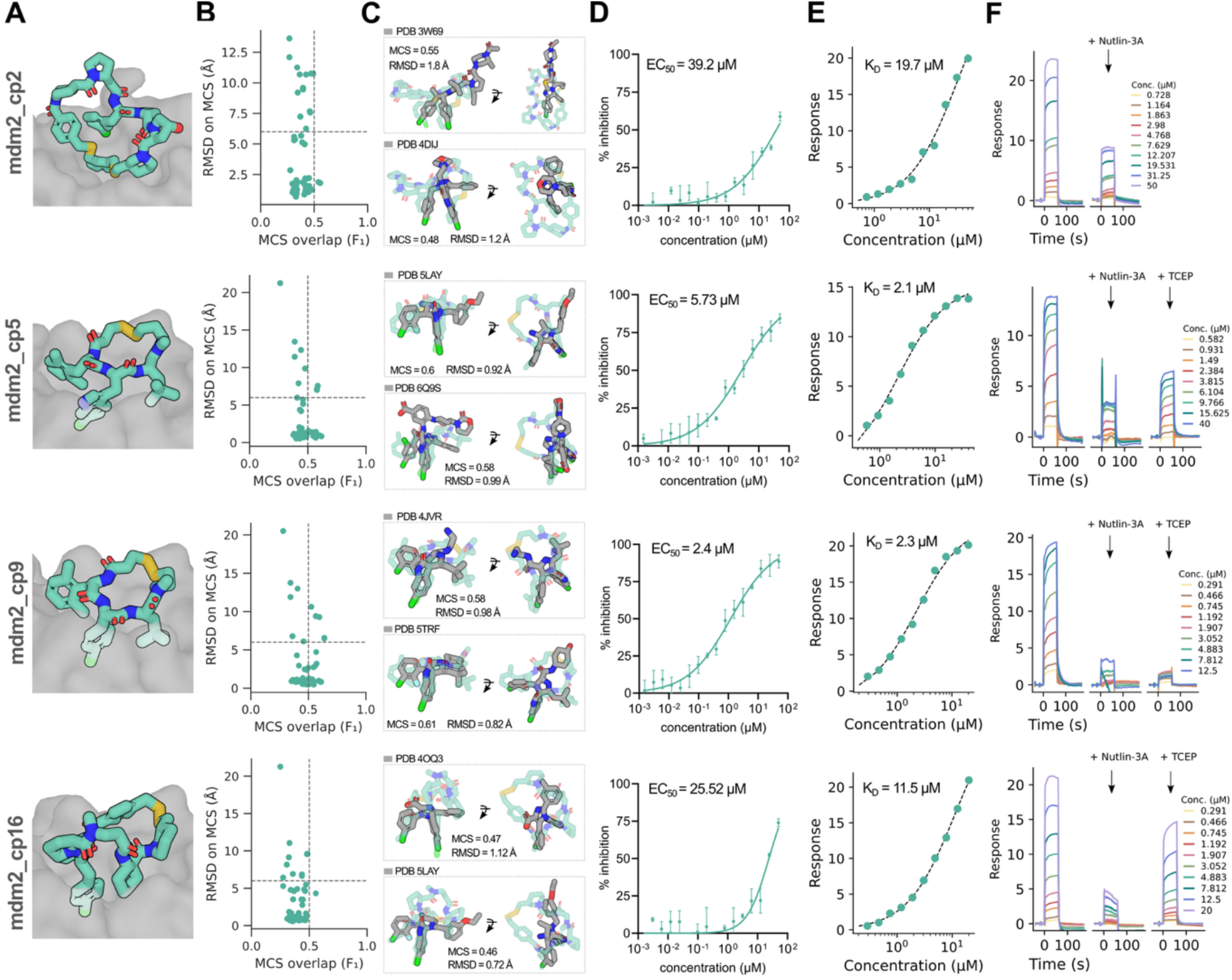
Computational and biochemical characterization for top *de novo* MDM2:p53 inhibitors. **A:** Structural models of designed cyclic peptide designs (colored and stick representation) bound to the canonical p53-binding site of MDM2 (shown as a gray surface). **B:** Structural similarity analysis comparing each cyclic peptide design to ligands observed in MDM2-bound structures from the PDB. Foldseek was used to identify relevant MDM2-ligand complexes from the PDB. Retrieved structures were aligned through the MDM2 protein, followed by MCS identification between each design and PDB ligand. The RMSD of the shared MCS fragment was then calculated. Scatter plots show MCS overlap fraction (reported as the F1 score) on the x-axis and MCS-fragment RMSD on the y-axis. Points with high MCS overlap and low RMSD represent ligands with the closest structural and spatial similarity to the designed cyclic peptides. **C:** Examples of PDB ligands exhibiting high MCS overlap and low RMSD relative to the cyclic peptide designs. **D:** Dose-response curves from the MDM2:p53 HTRF competition assay using Nutlin-3A competition, showing concentration-dependent disruption of the MDM2:p53 interaction by the cyclic peptide designs. Apparent EC_50_ values are indicated. **E:** Equilibrium SPR binding analysis showing concentration-dependent binding of cyclic peptide designs to MDM2, with fitted dissociation constants (K_D_) reported. **F:** Competitive SPR sensorgrams acquired in the presence and absence of the orthosteric MDM2 inhibitor Nutlin-3A. Reduced peptide binding in the presence of Nutlin-3A indicates competition for the canonical p53-binding pocket on MDM2 and supports binding of the designs to the same or an overlapping site. For disulfide-cyclized peptides, binding was measured in the presence of TCEP, which reduces the disulfide and is expected to generate the corresponding linear peptide.

We then asked whether the predicted MDM2-binding modes recapitulated interaction motifs observed in experimentally determined MDM2–ligand complexes (Fig 5A; Sup. Fig. 6.1-6.3A). To compare the structural models of our designs with known MDM2 ligands, we identified structurally related MDM2 complexes from the PDB, aligned the PDB structures to the predicted design complexes through the MDM2 protein, and calculated the MCS overlap and RMSD between PDB ligands and the designed cyclic peptides as done for Thrombin (Methods). The best-matching designs showed partial ligand overlap of approximately 50–55% with substructure RMSD values below 2 Å (Fig. 5B).

However, the full combination of the main MDM2-engaging ncAA motifs was not observed in any single analyzed PDB ligand. Instead, most matches involved one or two local building blocks (Fig. 5C; Sup. Fig. 13–16). This suggests that nCycle-Forge combined known hydrophobic interaction motifs into new cyclic peptide arrangements.

Two potent designs, mdm2_cp5 and mdm2_cp9, were structurally related, sharing three building blocks but differing in the central pocket-engaging motif. In the predicted binding models, mdm2_cp5 engages MDM2 through a chlorotryptophan derived motif, whereas mdm2_cp9 uses a chlorophenyl-containing motif in a similar region of the binding cleft (Fig. 5A–C; Sup. Fig. 14,15). Thus, closely related cyclic peptide scaffolds can support different hydrophobic ncAA motifs while retaining productive MDM2:p53 inhibition.

To determine whether functional inhibition reflected direct target engagement, we measured binding of purified MDM2 inhibitors to immobilized MDM2 by SPR. Compound concentration ranges were adjusted based on 1H NMR-derived effective solubility values (Methods; Sup. Table 2). Several compounds with measurable HTRF activity also showed concentration-dependent binding to MDM2 by SPR (Sup. Table 2). The two most potent designs, mdm2_cp5 and mdm2_cp9, bound MDM2 with apparent K_D_ values of 2.1 μM and 2.3 μM, respectively (Fig. 5E; Sup. Fig. 6.1–6.3F,G; Sup. Table 2). For comparison, Nutlin-3A bound with an apparent K_D_ of 0.04 μM, whereas the reference cyclic peptide M6 bound with an apparent K_D_ of 10.8 μM under the same conditions (Sup. Table 2; Sup. Fig. 6.3F,G). These data support that the designed cyclic peptides disrupt the MDM2 interaction by directly binding MDM2.

We next performed competitive SPR experiments with Nutlin-3A to test whether the designed peptides bind the canonical p53-binding cleft of MDM2. For compounds that showed measurable SPR binding, responses were reduced or abolished in the presence of Nutlin-3A, indicating competition with Nutlin-3A and supporting binding to the same or an overlapping site within the MDM2 p53-binding pocket (Fig. 5F; Sup. Fig. 6.1–6.3H).

Because p53 binds MDM2 as a short, structured α-helical linear peptide, we also asked whether the cyclic constraint was important for binding by testing disulfide-cyclized designs under reducing conditions. For these compounds, SPR measurements were performed in the presence of TCEP, which is expected to generate the corresponding linear peptide forms (Methods). Under these conditions, MDM2 binding signal was reduced or abolished for mdm2_cp5, mdm2_cp9, mdm2_cp16, and the reference peptide M6, whereas Nutlin-3A, which lacks a disulfide, was unaffected (Sup. Fig. 6.1–6.3I,J; Sup. Table 2). Notably, mdm2_cp9 lost detectable binding, while mdm2_cp16 retained weaker but detectable binding in the presence of TCEP (Fig. 5F; Sup. Table 2). These results suggest that cyclization and the resulting conformational constraint are important for productive MDM2 engagement.

Together, the HTRF inhibition data, direct SPR binding, Nutlin-3A competition, and TCEP linearization experiments support that nCycle-Forge can design cyclic peptides that disrupt a broad PPI interface such as the MDM2 binding site. The results further suggest that the method can arrange multiple hydrophobic ncAA motifs in a productive cyclic scaffold, where conformational constraint contributes to binding and function.

## Discussion

The computational *de novo* design of functional cyclic peptides incorporating ncAAs, diverse linker chemistries and cyclization formats remains a formidable challenge.

Here we describe nCycle-Forge, a computational framework that uses cofolding predictions as a surrogate fitness function to search the cyclic peptide chemical and structural space. nCycle-Forge combines gradient-free simulated annealing with atom-level representations of complete cyclic peptides, allowing building blocks and linkers to be varied without building block specific residue parameterization, precomputed conformer ensembles or model retraining.

Experimental validation of nCycle-Forge suggests that cofolding confidence, ipTM and ligand pLDDT provide useful, although imperfect, surrogate signals for cyclic peptide binder design. Inhibition of Thrombin activity and disruption of the MDM2:p53 interaction by nCycle-Forge-designed cyclic peptides indicate that the method readily enriches for functional molecules. This is encouraging because the two targets present distinct design challenges i.e., Thrombin contains a geometrically constrained enzymatic pocket, whereas MDM2 presents a broader PPI surface. The design of active compounds against both targets therefore supports the use of cofolding-derived scores for hit finding and library enrichment.

Consistent with this, the designed library yielded an approximately 6- to 24-fold higher hit rate against Thrombin than larger, target-agnostic crude libraries, while achieving a hit rate against MDM2 comparable to that of a target-tailored, hotspot-centered combinatorial library.

Other deep-learning methods have reported designed MDM2-binding macrocycles with comparable affinities in the low-micromolar range^22,34,35^, compared with 2.1 and 2.3 μM for mdm2_cp5 and mdm2_cp9. These used larger cyclic peptides built from the 20 canonical amino acids and recapitulated the native p53 interaction motif, presenting Phe, Trp and either Leu or Met from a helical segment. Across both comparisons, the designs reported here were obtained from the target sequence alone, guided primarily by cofolding scores, without fixing known binding motifs or exhaustively enumerating the combinatorial design space, reaching micromolar affinity directly through chlorinated ncAA motifs in smaller, non-helical scaffolds.

Although nCycle-Forge still requires screening hundreds of designs per target rather than a handful, these hits were obtained in one rapid design–synthesis–test round, rather than through the prolonged iterative cycles typically required for hit identification. The resulting hits were chemically and structurally diverse, spanning different cyclic peptide lengths, molecular sizes, and combinations of ncAA chemotypes. In many cases, the identified cyclic peptide inhibitors differed substantially from known target ligands in topology, size, and overall chemical composition. This suggests that the search does not simply converge to a single motif family or a known ligand and provides multiple starting points for further optimization. This differs from combinatorial screening approaches, for example against the same MDM2 interaction, where confirmed hits are often concentrated around a privileged scaffold and binding motifs^13,65,66^.

nCycle-Forge repeatedly enriched building blocks containing chemotypes known from ligands of the corresponding targets, including chlorothiophene and chlorophenyl moieties for Thrombin^13,14,67^, and indole moieties for MDM2^13,68^. Notably, these pharmacophores were recovered from the target sequence alone, without being explicitly rewarded or fixed during optimization. The same chemotypes were previously identified for these targets by synthesizing and assaying several-fold larger combinatorial libraries^13,14^, indicating that the search recovered established pocket-engaging motifs at a small fraction of the experimental cost. Because Thrombin- and MDM2-bound complexes are present in the PDB and likely represented in the training data, part of this enrichment may reflect learned priors over known chemotypes and binding geometries rather than generalization^69,70^, particularly for Thrombin where the higher hit rate coincided with enrichment of motifs that also dominated previous library hits.

A subset of confirmed hits however contained chemotypes without obvious precedent in experimentally solved target–ligand structures. The most notable example is thbn_cp22a, which positions an aza-tryptophan in the Thrombin active-site pocket.

Among the Thrombin-bound PDB ligands retrieved by our substructure analysis, none contained an equivalent moiety with comparable overlap and RMSD (Sup. Fig. 12). This design inhibited Thrombin with a K_i_ of 4.6 μM and bound with a K_D_ of 11 μM (Sup. Table 1).

Competition SPR supports binding at the intended sites, while TCEP treatment weakened the affinity of potent disulfide-cyclized designs, indicating that cyclization is important for activity. However, neither experiment establishes the predicted atomic binding mode. Experimental structures of representative peptide–target complexes therefore remain an important next step, particularly for validating the proposed novel pocket-engaging motifs.

Together, this work demonstrates that simulated-annealing searches guided by cofolding models can be integrated with automated synthesis and direct screening to identify functional ncAA-containing cyclic peptides. The modularity of nCycle-Forge should make it useful across a wide range of peptide design problems, including scaffolding known interaction motifs from larger molecules, such as antibodies or native ligands, into cyclic peptide formats, fully *de novo* cyclic peptide design, and co-optimization with objectives related to physicochemical properties or permeability. The same formulation could also enable explicit negative design by penalizing predicted engagement of homologous proteins or other off-targets, thereby supporting the simultaneous optimization of potency, selectivity and developability.

We anticipate that nCycle-Forge will be broadly useful for developing ncAA-containing cyclic peptide binders against challenging molecular targets across the proteome.

## Methods

### Estimating the theoretical accessible chemical space of small N-cyclic peptides

Linear peptides span a space of *B^n^* sequences for an *n*-mer drawn from an alphabet of size *B*. Here, we focus on the cyclic design space generated by installing a single linker between any pair of positions in an *n*-mer peptide. We assume *L* possible linkers and that the two linker attachment (“anchor”) positions must be chosen from a restricted set of linker-compatible building blocks of size *A*, while the remaining *n* − 2 positions are drawn from the full alphabet *B*. The number of cyclic candidates is therefore

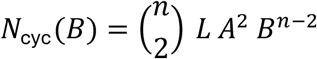

For example, for a 6-mer cyclic peptide *n* = 6, and using *B* = 1000 ncAAs, *L* = 40, and *A* = 25 yields

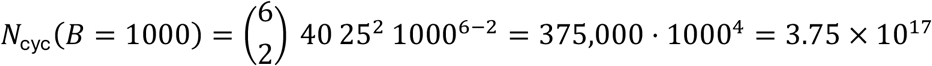

Taking into account small cyclic peptides of lengths 3-8 positions, the total number of cyclic candidates is over each length:

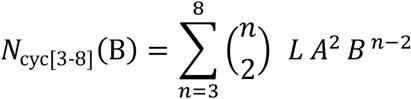

Using again *B* = 1000, *L* = 40, and *A* = 25, this evaluates to

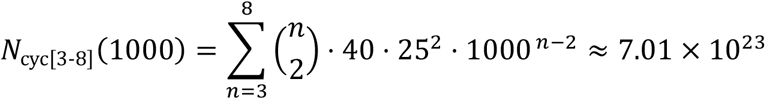

### Simulated annealing on a surrogate ligand–target interaction fitness landscape

Designing cyclic peptides to interact with targets can be viewed as navigating a ligand-target fitness landscape, a mapping that quantifies the fitness of a ligand to bind a specified surface region (i.e., site) on a target protein. We denote the true unknown interaction fitness landscape by *F*(*x*), where *x* is a ligand candidate. Importantly, this landscape is not directly accessible and would require experimentally testing many if not all possible ligands, which is often infeasible in practice.

Cyclic peptides were represented as sequences of discrete building blocks, including ncAAs and linkers. We denote the space of chemically admissible cyclic peptide sequences under the allowed building block alphabet and cyclization constraints as X, and let *x* ∈ X be a candidate cyclic peptide ligand. To explore X, we applied local perturbations at the building block level e.g., substitution of a single ncAA or linker per step. Thus, candidate proposals were sampled as

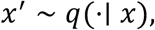

where *q* is a proposal distribution concentrated on a local neighborhood of *x*, thereby applying incremental sequence-level edits.

For each candidate ligand *x*, we performed a target-ligand cofolding step using Boltz to obtain a predicted complex

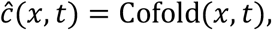

where *t* denotes the fixed target protein sequence context. From the predicted outputs which include several *in silico* scores, we defined a surrogate fitness function *F̂*(*x*), where higher values define a better predicted interaction fitness:

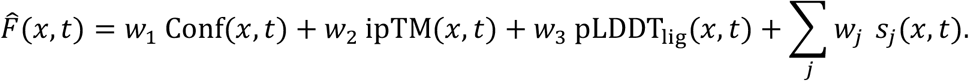

Here, Conf(*x*, *t*) is the model confidence term, ipTM(*x*, *t*) is the interface predicted TM-score, and pLDDT_lig_(*x*, *t*) is the ligand-local confidence (e.g., averaged across ligand atoms). The additional scoring terms *s*_j_(*x*, *t*) are optional and capture further design-relevant scores or penalties such as site engagement constraints, geometric plausibility filters, rotatable bond penalties, or charge-related terms. All weights {*w_i_*} were specified *a priori* and held fixed within an optimization trajectory.

We do not assume that the surrogate fitness *F̂* approximates the true interaction fitness *F* pointwise. Instead, we hypothesize that some local optima of *F̂* lie near local optima of *F* in the chemically admissible sequence space X. That is, for some surrogate optimum *x̂*\*, there exists a true optimum *d*(*x̂*\*, *x*\*) ≤ *ε*, where *d* measures the edit distance. Under this hypothesis, optimizing *F̂* can guide the search toward regions near true interaction peaks. We do not expect this correspondence to hold for every surrogate optimum, and some trajectories may therefore converge to spurious high-scoring solutions.

We optimized ligands using simulated annealing on the surrogate landscape *F̂*: *X* → ℝ. Starting from a randomly initialized cyclic peptide *x*_0_, at iteration *k* we:

1. Proposed a local perturbation *x*^′^ ∼ *q*(⋅∣ *x_k_*),
2. Evaluated *F̂*(*x*′, *t*) via cofolding,
3. Accepted or rejected the proposal according to a Metropolis acceptance rule:

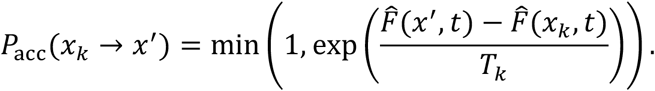

The temperature parameter *T_k_* > 0 was exponentially annealed according to

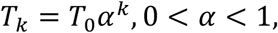

such that *T_k_* → 0 over the course of the design trajectory. At high temperature, the algorithm allows occasional downhill moves on *F̂*, enabling exploration of sequence space and escape from local optima. As *T_k_* decreases, the algorithm progressively approaches greedy hill-climbing on the surrogate landscape.

The state was updated as:

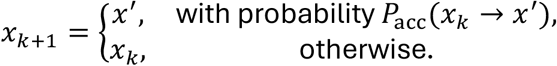

To overcome local optima traps, we ran many independent short trajectories of 100 steps each in parallel, each initialized from a different random starting ligand.

Although we used Boltz as the cofolding surrogate model in this work, the same simulated annealing framework is general and can theoretically be applied with any structure-based ligand–target cofolding or scoring method that provides a surrogate interaction objective.

### Co-optimization using guidance and penalties

#### Rule-of-5 penalty

To enable co-optimization of physicochemical properties alongside structural fitness (Fig. 1E), we incorporated a penalty term based on Rule-of-5 descriptors computed with RDKit (https://www.rdkit.org/). For each cyclic peptide, we evaluated hydrogen-bond donors (HBD), hydrogen-bond acceptors (HBA), molecular weight (MW), Wildman–Crippen log P, topological polar surface area (TPSA), and number of rotatable bonds, with upper-bound thresholds of HBD ≤ 5, HBA ≤ 10, MW ≤ 500 Da, log P ≤ 5, TPSA ≤ 140 Å², and ≤ 10 rotatable bonds. Descriptors were min-max normalized to [0,1] using fixed global ranges. For each violated constraint, we defined a normalized violation *δ* as the difference between the scaled value and the scaled threshold. Violations were transformed into non-negative penalties via 10^ẟ^ − 1 and summed across properties. The resulting score,

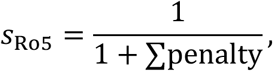

equals 1 when all constraints are satisfied and decreases with increasing violations. The term was added to the surrogate objective, enabling trajectories to simultaneously improve structure-based scores while reducing Ro5 violations, as shown in Fig. 1E.

#### Motif integration via contact guidance restraints

To scaffold known interaction motifs into cyclic peptides (Fig. 1F), we derive contact guidance restraints from a reference complex containing the receptor and a motif in the desired binding geometry. Receptor– motif atom pairs within a cutoff (typically 6 Å) are identified and their corresponding distances are recorded. To transfer these interactions onto the designed ligand, motif atoms are mapped onto ligand atoms via chemical correspondence (substructure or maximum common substructure matching). Each motif specification defines a ligand building block index in the cyclic peptide sequence, and only pairs involving atoms at this position are kept. For each retained pair, a distance restraint is defined using the corresponding reference separation. These restraints are provided to the structure model during cofolding, biasing sampling toward conformations that preserve the motif geometry while allowing the surrounding cyclic scaffold to adapt.

#### Hotspot diversification via contact guidance restraints

To enable exploration of chemically diverse substituents while preserving key interaction geometry (Fig. 1G), we implemented hotspot diversification using the same contact guidance framework as described above. A reference motif capturing the desired receptor contacts was used to derive distance restraints. Motif atoms were mapped onto ligand atoms via substructure or maximum common substructure matching. Because motif and building block substituents can differ chemically, only atom pairs that could be mapped between the motif and the sampled building block were retained. The motif (hotspot) was anchored to a specific residue position in the cyclic peptide, while allowing alternative building blocks to be sampled at that position. Other sequence positions were varied independently, enabling diversification of the cyclic scaffold while maintaining the hotspot geometry. As above, these constraints act through the structure model’s internal guidance potentials and are not included as an explicit scalar term in the optimization objective.

### Scoring, filtering and selection of cyclic peptide designs

#### Scoring and filtering with Rosetta

To evaluate the physicochemical quality of predicted cyclic peptide-receptor complexes, each design was scored using Rosetta^71^. First, Boltz-predicted complexes were preprocessed by separating ligand and receptor chains, removing water and non-proteinogenic residues from the receptor, and adding explicit hydrogens to the ligand using OpenBabel^72^ at physiological pH (7.4). Ligand topology and parameter files were generated using Rosetta’s “molfile_to_params” script. The protonated complex was then energy-minimized using a Cartesian MinMover under the ref2015 score function (with cart_bonded weight 0.625 and pro_close set to 0.0), allowing receptor side-chain conformations and bond geometry to relax while keeping ligand coordinates fixed. The minimized complex (after recomputing topology parameters) was used to compute interface properties using the InterfaceScoreCalculator and InterfaceAnalyzerMover, together with a ShapeComplementarity filter and LigInterfaceEnergy filter. The resulting per-design metrics included dG_separated, dSASA_int, delta_unsatHbonds, hbonds_int, ligand_e_iface, shape complementarity (sc), and component energy terms (fa_atr, fa_rep, fa_sol).

#### Design selection

The library of 1,472 cyclic peptides was assembled in two stages. In the first stage, the design pool was filtered to retain candidates satisfying confidence_score > 0.8, ipTM > 0.8, ligand pLDDT > 0.8, ligand self-consistency RMSD < 2.5 Å (an RMSD between two independent cofolding runs, one potentially constrained and one unconstrained), and Rosetta dG_separated < 0. To limit over-representation of problematic building blocks, per building block caps were applied based on objective score. In the second stage, remaining slots were filled with designs drawn from the full trajectory outputs, requiring confidence_score, ipTM, and ligand pLDDT all > 0.85 and excluding sequences already selected. Candidates were ranked by the sum of confidence_score, ipTM, and ligand pLDDT, and accepted using a Jaccard-distance diversity cutoff of 0.8 computed over the set of ncAAs. Additional per building block frequency caps were applied as above.

### Chemical diversity and substructure comparison analyses

#### Chemical diversity comparison to ChEMBL and reference cyclic peptide libraries

Chemical diversity of nCycle-Forge designs was compared to target-specific ChEMBL compounds and previously reported cyclic peptide libraries from Habeshian et al.^13^. For each target, we compared four compound sets: the deduplicated nCycle-Forge design pool, the subset selected for synthesis, the corresponding Habeshian et al. enumerated cyclic peptide library, and target-associated ChEMBL compounds. ChEMBL reference compounds were obtained from target-specific ChEMBL activity exports for human Thrombin/prothrombin (Target ChEMBL ID: CHEMBL204) and human MDM2 (Target ChEMBL ID: CHEMBL5023). For each target, activity records were restricted to ChEMBL binding assays (K_i_ or IC_50_). To avoid repeated representation of the same molecule, only one activity record was retained per ChEMBL ID. Compounds were then filtered to retain molecules with reported K_i_ or IC_50_ values ≤ 1000 nM. Compounds with molecular weight <100 Da or >2500 Da were removed prior to chemical standardization and fingerprint calculation. All molecules were standardized using RDKit MolStandardize by normalizing functional groups, reionizing, selecting the largest parent fragment, uncharging, and generating the canonical Simplified Molecular Input Line Entry System (SMILES). Molecules that could not be parsed or standardized were excluded. For each molecule, MinHash fingerprints (MHFP) were generated using 2,048 permutations with radius 1. Pairwise fingerprint distances were computed and embedded into two dimensions using t-SNE as implemented in openTSNE (https://opentsne.readthedocs.io/), with perplexity 256, learning rate 1,000, early exaggeration 48, and 2,000 iterations. Thrombin and MDM2 were embedded separately. The resulting embeddings were used only for visualization of chemical-space coverage and relative overlap, not for quantitative distance comparisons.

#### Comparison of designed cyclic peptides to PDB ligands at the ncAA-fragment level

To assess whether designed cyclic peptides contained motifs related to ligands observed in experimentally solved structures, we compared selected Thrombin and MDM2 designs to small molecule ligands in structurally related PDB complexes. For each target, a representative predicted protein–cyclic peptide complex was used as a Fold-seek^73^ query. The protein chain was extracted from the predicted complex and searched against the PDB100 database using Foldseek. Hits were retained if they had a TM-score ≥ 0.5. Duplicate PDB IDs were removed by retaining the highest-scoring Foldseek hit. For each retained PDB entry, the primary non-polymeric ligand was extracted from the corresponding processed structure file, requiring at least five heavy atoms. Ligands were then reconstructed as RDKit molecules.

A whole-ligand maximum common substructure (MCS) was calculated using RDKit. For the global MCS, the fraction of the design and reference ligand covered by the MCS was recorded, and the harmonic mean of the two fractions was used as an MCS-overlap F1 score. To quantify spatial similarity, the heavy-atom RMSD over the whole-ligand MCS was calculated after alignment on the target protein. When multiple symmetry-equivalent MCS mappings were possible, all mappings were enumerated where feasible, and the minimum heavy-atom RMSD was retained.

For ncAA-level comparisons, each building block was compared against each transformed PDB ligand using an RDKit MCS search. MCS matching required matching atom elements and bond orders, ring atoms to match ring atoms, and complete rings to be preserved. For each ncAA–PDB-ligand MCS, all symmetry-equivalent substructure mappings were enumerated where possible, and the minimum heavy-atom RMSD over the mapped atoms was retained. For each match, we recorded the ncAA fragment identity, the corresponding substructure-overlap F1 score, and the RMSD of the matched atoms after protein-based alignment. Matches were ranked by increasing substructure RMSD and decreasing overlap score. Representative examples were visualized by highlighting the matched ncAA fragment in the designed cyclic peptide and the corresponding MCS in the PDB ligand.

### Automated peptide library synthesis in 384-well plates

Automated solid-phase peptide synthesis was performed on a CEM Multipep 2 synthesizer that has been modified to accept a capacity of four 384-well plates as previously described^74^. Disulfide linker resins were prepared as previously described^75^ and roughly 2.5 μmoles were transferred to each well of a 384-well solid phase synthesis plate (Agilent 201035-100).

The resin in each well was washed with 1 × 50 μL DMF. Coupling was performed first with 25 μL of Fmoc amino acid solution (500 mM, 5 equiv.), 13 μL DIC solution (1 M, 5.2 equiv.), 8 μL of Oxyma solution (1 M, 3.2 equiv.), and 5 μL *N*-methylpyrrolidone. All components were premixed for 1 min, then added to the resin for 2 h reaction without shaking. The final volume of the coupling reaction was 51 μL and the final concentrations of reagents were 245 mM amino acid, 255 mM DIC, and 157 mM Oxyma. The resin was then washed with 2 × 65 μL of DMF. A second coupling was performed with 26 μL of Fmoc amino acid solution (500 mM, 5.2 equiv.), 25 μL HATU solution (500 mM, 5 equiv.), 7 μL of *N*-methylmorpholine solution (4 M, 11.2 equiv.), and 5 μL *N*-methylpyrrolidone. All components were premixed for 1 min, then added to the resin for 2 h reaction without shaking. The final volume of the coupling reaction was 63 μL and the final concentrations of reagents were 206 mM amino acid, 198 mM HATU and 444 mM *N*-methylmorpholine. The resin was then washed with 2 × 65 μL of DMF. Fmoc deprotection was performed using 50 μL of 2% DBU in DMF (v/v) for 2 min, followed by washing with 6 × 65 μL DMF.

### Library deprotection and cleavage

Sequences which did not contain side chain protection were not subjected to deprotection conditions. For those with protecting groups, 60 μL of deprotection mixture (95% TFA, 2.5% TIPS, 2.5% H_2_O (v/v/v)) was added using a multichannel pipette and allowed to incubate for 30 minutes. The TFA solution was allowed to drip through the filter plate into a collection plate during this time. The treatment was repeated once more, and the resin in each well was washed with 5 × 80 μL of DCM and subsequently allowed to air dry for 30 min.

Reductive cleavage was performed by treatment twice with 30 μL of DMF containing 333 mM 1,4-butanedithiol (BDT) and 667 mM triethylamine (TEA), first with 2 hours incubation and then repeated with overnight incubation. The cleavage solution was dispensed using a specially modified Certus Flex liquid dispenser (Fritz Gyger AG). After each incubation, DMF solution was collected into a 384-well receiver plate (Labcyte PP-0200) by centrifugation at 395 RCF (Sigma 4-16KS) for 2 min.

To the recovered DMF stock was added 5 uL of formic acid in milliQ-water (40% (v/v), 1.3 equiv. relative to TEA), and the peptides were dried by RVC (30-40 °C, 1500 rpm, 0.1 mbar). Once dry (∼3 hours), the peptides were redissolved in 25 μL DMSO for further use.

### Library peptide recovery quantification by absorption

Ellman’s reagent (5,5′-dithiobis-(2-nitrobenzoic acid)) was dissolved in assay buffer (60 mM NH_4_HCO_3_ in 1:1 water/MeCN, pH 8 to a concentration of 10 mM. To a 384-well plate with transparent bottom (Greiner 781162) was transferred dithiol peptide in DMSO (60 nL) by acoustic droplet ejection (Beckman Coulter Echo 650). Then, 25 µL of the Ellman’s reagent solution was dispensed to each well, and absorbance at 412 nm was measured on a BMG Labtech Pherastar FSX plate reader. Peptide concentrations were calculated based on a calibration curve of beta-mercaptoethanol using the below equation:

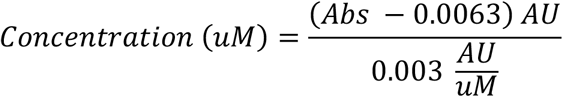

where Abs is the absorption (AU) at 412 nm as measured on the plate reader.

### Concentration adjustment and transfer to cyclization plates

According to the results of the Ellman’s assay, additional DMSO was added to the plates containing linear peptide stocks to bring each plate to a uniform average concentration. Using a 384-channel pipettor (Integra Viaflo 384), a copy of each plate was made containing on average 50 nmoles of peptide in each well. The plates were then dried by RVC (30**-**40 °C, 1500 rpm, 0.1 mbar) for 1 hour.

| Plate | Average concentration | Volume to transfer (50 nmoles peptide) |
| --- | --- | --- |
| Plate 1 | 7 | 7.14 |
| Plate 2 | 8 | 6.25 |
| Plate 3 | 12 | 4.17 |
| Plate 4 | 5 | 10 |

### Re-reduction and library cyclization

The linear peptides at 50 nmole scale were re-reduced to ensure no disulfide from spontaneous oxidation was present prior to cyclization. To each well was added 16 μL of 12.5 mM BDT and 100 mM TEA in DMF and reduction was allowed to proceed for 1 hour. After this time, 4 μL of 2% (v/v) formic acid in H_2_O (1.3 equiv. relative to base) was added, and the plates were dried by RVC (30-40 °C, 1500 rpm, 0.1 mbar) for 2 hours.

Bis-electrophilic linkers were dissolved in a 100 mM solution of ammonium bicarbonate in 1:1 H_2_O/MeCN at pH 8.0 to a concentration of 3 mM. The appropriate linker solutions (50 μL, 3 equiv. relative to peptide) were dispensed to wells using a Certux Flex liquid dispenser and used to dissolve the reduced linear di-thiol peptide in each tube to a final peptide concentration of 1 mM. The plates were allowed to incubate for 2 hours at room temperature, after which time excess linker was quenched with 5 μL of a 120 mM β-mercaptoethanol solution in H_2_O (4 equiv. relative to linker) overnight. The next day, the plates were dried by RVC (30-40 °C, 1500 rpm, 0.1 mbar).

Cyclic disulfides were synthesized by dissolving dry linear peptides in 20% DMSO, 30% MeCN, and 50% 200 mM ammonium bicarbonate buffer pH 8.0 (v/v/v) and allowing to oxidize for 2 days at room temperature. The plates were then dried by RVC (30-40 °C, 1500 rpm, 0.1 mbar).

### LC-MS analysis

Peptides were analyzed by LC-MS analysis with a UHPLC (Shimadzu Nexera LC40) and single quadrupole MS system (Shimadzu LCMS-2020) using a C18 reversed phase column (Phenomenex Kinetex 2.1 mm × 50 mm C18 column, 100 Å pore, 2.6 μm particle) and a linear gradient of solvent B (acetonitrile, 0.05% formic acid) over solvent A (H_2_O, 0.05% formic acid) at a flow rate of 0.6 mL/min. Mass analysis was performed in positive ion mode. Data was collected and analyzed using Shimadzu LabSolutions software.

For the LC-MS analysis, the samples of the various experiments were prepared as follows. Linear or cyclic peptide stocks were diluted into 1:1 water/acetonitrile + 0.1% formic acid to give a peptide concentration of 200 μM. For all analyses, 10 μL of the samples were injected, typically using a 0 to 100% gradient of solvent B over 5 min.

### mg-scale synthesis and purification of Thrombin and MDM2 hits

Selected Thrombin and MDM2 hits were re-synthesized and purified on a compound-by-compound basis either in-house or by WuXi AppTec.

For in-house syntheses, peptides were synthesized on a CEM Multipep 2 automated solid-phase peptide synthesis at a 50 μmole scale using disposable 5 mL syringes. Resin was added to each syringe and was washed with 3 × 2 mL DMF. Coupling was performed first with 500 μL of Fmoc amino acid solution (500 mM, 5 equiv.), 260 μL DIC solution (1 M, 5.2 equiv.), 160 μL of Oxyma solution (1 M, 3.2 equiv.), and 100 μL *N*-methylpyrrolidone. All components were premixed for 1 min, then added to the resin for 2 h reaction with shaking. The final volume of the coupling reaction was 1020 μL and the final concentrations of reagents were 245 mM amino acid, 255 mM DIC, and 157 mM Oxyma. The resin was then washed with 2 × 2 mL of DMF. A second coupling was performed with 520 μL of Fmoc amino acid solution (500 mM, 5.2 equiv.), 500 μL HATU solution (500 mM, 5 equiv.), 140 μL of *N*-methylmorpholine solution (4 M, 11.2 equiv.), and 100 μL *N*-methylpyrrolidone. All components were premixed for 1 min, then added to the resin for 2 h reaction with shaking. The final volume of the coupling reaction was 1260 μL and the final concentrations of reagents were 206 mM amino acid, 198 mM HATU and 444 mM *N*-methylmorpholine. The resin was then washed with 2 × 2 mL of DMF. Fmoc deprotection was performed using 1 mL of 2% DBU in DMF (v/v) for 1 min, followed by washing with 6 × 2 mL DMF.

Peptides were deprotected by different solutions according to the specific protecting groups present to minimize exposure to high percentages of TFA:

| Solution | Protecting groups |
| --- | --- |
| 20% TFA, 2.5% H<sub>2</sub>O, 2.5% TIS, 75% DCM (v/v/v/v) | Trityl |
| 50% TFA, 2.5% H<sub>2</sub>O, 2.5% TIS, 45% DCM (v/v/v/v) | Boc |
| 95% TFA, 2.5% H<sub>2</sub>O, 2.5% TIS (v/v/v) | tBu ethers |

In all cases, the resin was incubated with 3 mL of deprotection mixture for 2 ×15 min on an orbital shaker. The deprotection solutions were discarded, and the resin washed with 3 × 3 mL DCM.

Reductive cleavage was performed by treatment twice with 3 mL of DMF containing 70 mM 1,4-butanedithiol (BDT) and 130 mM triethylamine (TEA), first with 1 hour incubation and shaking, and then repeated with overnight incubation and shaking. The DMF solution was collected into a 50 mL conical tube after each treatment. To the recovered DMF stock was added 200 μL of TFA in milliQ-water (40% (v/v), 1.28 equiv. relative to TEA), and the peptides were dried by RVC (30-45 °C, 1500 rpm, 0.1 mbar).

### Cyclization

Bis-electrophilic linkers were dissolved in a 100 mM solution of ammonium bicarbonate in 1:1 H_2_O/MeCN at pH 8.0 to a concentration of 3 mM. For scaled peptide synthesis, no quantification following cleavage was performed, and recovery was estimated to be 50%. The appropriate linker solutions (25 mL, 3 equiv. relative to peptide) were used to dissolve the reduced linear di-thiol peptide in each tube to a final peptide concentration of 1 mM. The tubes were shaken for 2 hours at room temperature, after which time excess linker was quenched with neat β-mercaptoethanol (42 μL, 8 equiv. relative to linker) for 2 hours. Tubes were then lyophilized to dryness.

Cyclic disulfides were synthesized by dissolving dry linear peptide in 20% DMSO, 30% MeCN, and 50% 200 mM ammonium bicarbonate buffer pH 8.0 (v/v/v) and allowing to oxidize for 2 days at room temperature.

### Purification

Compounds synthesized in-house were purified by RP-HPLC using a Shimadzu HPLC system equipped with an SPD-40 UV detector, LC-20AP pumps, and an LH40 fraction collector, using a 19 mm × 250 mm Waters XBridge C18 OBD preparative column (130 Å pore size, 5 μm particle size). Solvent A was H_2_O with 0.1% formic acid (v/v), and solvent B was MeCN with 0.1% formic acid (v/v). Purified compounds from both in-house and WuXi AppTec syntheses were used for downstream assays after confirmation of the expected mass by LC-MS and assessment of chromatographic purity by analytical HPLC or UPLC. For compounds purified by WuXi AppTec, analytical purity was assessed by HPLC/UPLC UV absorbance at 220 nm using vendor-provided analytical gradients, and molecular-weight confirmation was obtained from LC-MS chromatograms and corresponding MS spectra. Reported purity values were derived from UV chromatographic peak integration.

### Thrombin inhibition assay

Thrombin activity in presence of macrocycles and reference compounds was assessed using a fluorogenic substrate turnover assay^13,14^. Compounds dissolved in DMSO were transferred to 384-well assay plates using a Labcyte Echo 650 acoustic dispenser. Compounds from the crude library were screened at a final concentration of 10 μM. In dose-response curves with pure compounds, 16 two-fold dilutions were tested, starting from 50 μM highest concentration. Thrombin purified from human plasma at a final concentration of 2 nM in Tris buffer at pH 7.4 (100 mM Tris-Cl, 150 mM NaCl, 10 mM MgCl2, 1 mM CaCl2, 0.1% w/v BSA, 0.01% v/v Triton-X100) was dispensed into each well using a Gyger Certus Flex liquid dispenser and incubated for 10 min at room temperature. The fluorogenic substrate Z-Gly-Gly-Arg-AMC at a final concentration of 25 μM in the same Tris buffer was added to each well using the liquid dispenser. Florescence intensity was measured with a Hidex fluorescence plate reader (excitation at 355 nm, emission at 460 nm) at RT for a period of 45 min with a read every 5 min. The slope of fluorescence increase over time was calculated for each sample using Microsoft Excel. Values were subtracted for the negative control (DMSO, 0% inhibition) and the percentage of Thrombin inhibition was calculated dividing the slope of each sample by the slope of the 100% inhibition control (100 nM Dabigatran) and multiplying the results by 100. For the determination of the K_i_ values of the compounds, the percent of Thrombin inhibition was plotted against the corresponding compound concentrations, and sigmoidal curves were fitted using the following four-parameter equation in GraphPad Prism 10:

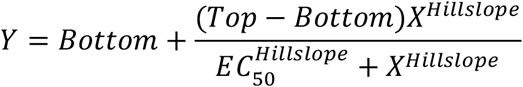

K_i_ values were determined from the EC_50_ values using the Cheng-Prusoff equation and K_m_ = 67 μM (determined for Thrombin and the fluorogenic substrate in the assay conditions):

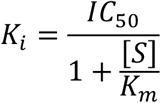

### MDM2 binding screen

MDM2 binding by cyclic peptides was assessed by measuring disruption of the interaction with p53 in a HTRF assay. Compounds dissolved in DMSO were transferred to 1536-well assay plates using a Labcyte Echo 650 acoustic dispenser. Compounds from the crude library were screened at a final concentration of 10 μM. In dose-response curves with pure compounds, 16 two-fold dilutions were tested, starting from 50 μM highest concentration. Human GST-tagged MDM2 (BPS) at a final concentration of 10 nM in 10 mM Hepes buffer at pH 7.4, 150 mM NaCl, 0.05% w/v BSA, 0.05% v/v Tween 20 was dispensed into each well using a Gyger Certus Flex liquid dispenser and incubated for 20 min at room temperature. Human FLAG-tagged p53 (BPS) in the same buffer was added to each well using the liquid dispenser at a final concentration of 5 nM. After 20- min incubation HTRF Anti-FLAG M2 Terbium-conjugate monoclonal antibody and Anti-GST d2-conjugate monoclonal antibody were prepared in the same buffer and added to each well using the liquid dispenser at a final 0.25x dilution. After 20-min incubation in the dark at room temperature, HTRF ratio was measured with a Pherastar reader (excitation at 337 nm, emission at 620 and 665 nm). Values were subtracted for the negative control (DMSO, 0% inhibition) and the percentage of MDM2:p53 PPI inhibition was calculated dividing for the HTRF ratio of the 100% inhibition control (50 μM Nutlin-3a) and multiplying the results by 100. For the determination of the EC_50_ values of the compounds, the percent of inhibition was plotted against the corresponding compound concentrations, and sigmoidal curves were fitted using the following four-parameter equation in GraphPad Prism 10:

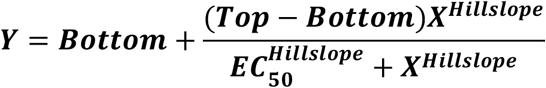

### Solubility determination

Compound solubility was assessed prior to SPR measurements and used to define the maximum compound concentrations used in subsequent SPR titrations.

Soluble compound concentrations were determined by quantitative one-dimensional ¹H NMR spectroscopy. For Thrombin compounds, test compounds were diluted from DMSO stocks to a nominal concentration of 100 μM in deuterated PBS pH 7.2 containing 5% d6-DMSO and 100 μM dTSP as an internal concentration standard. For MDM2 compounds, test compounds were prepared at nominal concentrations of up to 200 μM in deuterated buffer conditions matched as closely as possible to the cor-responding SPR assay buffers. These included 25 mM deuterated HEPES pH 7.4, 200 mM NaCl, 5% d6-DMSO, with or without 1 mM deuterated TCEP. For selected MDM2 compounds, additional pH conditions were assessed using deuterated Bi-Tris pH 6.4 or deuterated Tris pH 8.4.

¹H NMR spectra were acquired at room temperature on a 600 MHz spectrometer. Soluble compound concentrations were calculated by comparing the integrated intensity of resolved compound proton resonances with the dTSP internal standard. A 100 μM dTSP signal corresponds to 900 μM proton concentration and was used for quantification.

Compounds for which no assignable compound resonances were detected, or for which signals were too weak to integrate reliably, were reported as having solubility below the lower limit of detection of the assay, approximately 20 μM. The resulting effective solubility values were used to define the maximum compound concentrations used in subsequent SPR titrations.

### Surface plasmon resonance binding and competition assays

Compound concentration ranges were selected based on the solubility measurements described above. SPR sensorgrams were reference-subtracted and corrected using blank buffer injections where applicable. Equilibrium binding responses were fitted using steady-state a binding model, or sensorgrams were fitted to a 1:1 kinetic binding model using the Biacore Insight Evaluation software version 6 to determine KD values, unless otherwise indicated.

#### Thrombin SPR binding and competition assays

Thrombin SPR measurements were performed and analyzed on a Biacore T200 instrument at 20 °C using human α-Thrombin. Approximately 1,500 response units (RU) of human α-thrombin were immobilized on a Cytiva CM5 sensor chip by amine coupling, as described previously^76^. Thrombin surfaces were prepared using reconstituted Thrombin and, where indicated, SEC-repurified Thrombin. In some immobilizations, Melagatran was included to occupy the active site during coupling. The running buffer was 10 mM HEPES pH 7.5, 150 mM NaCl, 0.005% Tween-20, 3.4 mM EDTA, and 5% DMSO. Compounds were injected at 30 µl/min as 10-point, two-fold dilution series, with top concentrations adjusted according to compound solubility. Melagatran and Dabigatran were included as positive-control ligands to assess Thrombin surface activity. Generally, this immobilization procedure resulted in approximately 80% of the immobilized protein being active for control ligands binding. For competition experiments, compound bind-ing was measured in the absence and presence of 200 nM Melagatran to determine whether the designed cyclic peptides bound to the same or an overlapping active-site pocket.

#### MDM2 SPR binding and competition assays

MDM2 SPR measurements were performed on a Biacore 8K Plus instrument at 20 °C using human MDM2(17–125)-His6-AviTag immobilized on a Cytiva NTA Series S sensor chip by Ni-NTA capture coupling approach. Flow cell 1 was used as a reference channel and flow cell 2 contained immobilized MDM2, with an immobilization level of approximately 1,500 RU. The capture buffer was 25 mM HEPES pH 7.4, 200 mM NaCl, 1 mM TCEP, and 0.005% Tween P20. Binding measurements were performed in two running buffers: buffer A, consisting of 25 mM HEPES pH 7.4, 200 mM NaCl, 0.005% Tween P20, and 5% DMSO; and buffer B, consisting of the same buffer supplemented with 1 mM TCEP. Compounds were injected at 30 µl/min as 10-point concentration series with top concentrations adjusted according to NMR-derived solubility values. Nutlin-3A was included as a positive-control ligand and was titrated as a two-fold dilution series. Approximately 60% of immobilized protein was active for positive-control binding. For competition experiments, compound binding was measured in buffer A supplemented with 500 nM Nutlin-3A to assess competition with the canonical p53-binding pocket of MDM2.

## Acknowledgements

We thank Anna Belorusova, Johan Hollander, Denise Tsagris, Yevhenniia Nesterenko, Johan Veerman and Magdalena Ortiz at ZoBio for experimental support with compound solubility assessment, SPR assay execution and Thrombin protein preparation.

## Author contributions

O.H. and Z.H. conceived the study and designed the experiments. Z.H., F.R., T.B. and O.H. developed the software. Z.H. and F.R. performed the computational design and computational analysis of the library with support of T.B.. A.G. synthesized the crude cyclic peptide library with support from C.G. and S.H. C.G. performed the in-house re-synthesis and purification of selected cyclic peptide hits with support from A.G. and S.H. A.G., C.G. and S.H. performed chemical quality control and analyzed LC-MS and HPLC data for crude and purified compounds. C.T. established and performed the Thrombin enzymatic inhibition assay and the MDM2 HTRF assay and analyzed biochemical screening and dose-response data for crude and purified compounds. R.J.-B. and M.K. performed and coordinated compound solubility measurements and SPR experiments. R.J.-B., M.K. and G.S. analyzed and interpreted the solubility and SPR data. Z.H. prepared the figures and wrote the original draft of the manuscript. All authors analyzed data, reviewed and commented on the manuscript. O.H., S.H. and L.T. supervised the research.

## Competing interests

Z.H., F.R., C.T., A.G., C.G., T.B., S.H., L.T., and O.H. are current or former employees of Orbis Medicines, a cyclic peptide therapeutics company. R.J.-B., M.K., and G.S. are employees of ZoBio, which was contracted by Orbis Medicines to perform services related to the work described in this manuscript.

## Data availability

Code will be made available upon publication.

## Supplementary Methods

### Boltz cofolding test on a macrocyclic dataset

We used the macrocyclic dataset reported by Robson-Tull and Rodrigues^1^, which comprises 240 macrocycles in complex with their target proteins. For each entry, we downloaded the biological assembly and extracted all protein chains containing at least one atom within 6 Å of the macrocyclic ligand. The macrocyclic ligand was isolated and corrected using the chemical component dictionary (CCD)^2^.

Boltz input YAML files were prepared by encoding each protein chain as an AA sequence, while the macrocyclic ligand was provided as a SMILES string. Because our goal was to evaluate local cofolding accuracy, this is recovering the correct pose rather than identification of the binding site, we applied pocket constraints during cofolding predictions. To define the pocket, we selected the protein residue whose atoms were on average closest to the macrocyclic ligand in the experimental structure.

Boltz was run independently on each of the 240 complexes, and predictions were evaluated using (i) the RMSD over all ligand heavy atoms and (ii) an interface ligand RMSD, computed over ligand atoms located within 4 Å of the protein in the experimental structure.

## Supplementary Tables

**Sup. Table 1.** Thrombin inhibitors selected for re-synthesis and experimental characterization.

| Name | fraction | LCMS Purity (%) | EC50 (μM) | Ki (μM) | NMR solubility (max. effective concentration μM) | steady-state KD (μM) | k_on (1/Ms) | k_off (1/s) | kinetic KD (μM) | Competitive with Melagatran |
| --- | --- | --- | --- | --- | --- | --- | --- | --- | --- | --- |
| Melagatran |  |  |  |  |  | 0.0009 | 6.1 × 10<sup>7</sup> | 0.067 | 0.0011 | yes |
| Dabigatran |  |  | 0.00165±0.0008 | 0.0011±0.00058 |  | 0.0008 | 1.1 × 10<sup>7</sup> | 0.014 | 0.0013 | yes |
| M1 (ref) |  | 94.9 | 0.039±0.03 | 0.028±0.02 | 38 | 0.3 | 224000 | 0.0803 | 0.4 | yes |
| thbn_cp1 |  | 94 | 10.00±2.45 | 7.30±1.79 |  |  |  |  |  |  |
| thbn_cp2 |  | 94 | 1.33±0.55 | 0.97±0.40 | 72 | 9.3 |  |  |  | yes |
| thbn_cp3 |  | 95.2 | 0.61±0.05 | 0.43±0.03 | NA | 4.0 | 22800 | 0.09 | 3.9 | yes |
| thbn_cp4 |  | 96.85 | 1.89±0.30 | 1.34±0.21 | 56 | 12.6 |  |  |  | yes |
| thbn_cp5 |  | 77 | 0.32±0.16 | 0.23±0.12 |  |  |  |  |  |  |
| thbn_cp6 |  | 89 | 2.93±0.87 | 2.13±0.63 | 71 | 25.4 | 2280 | 0.042 | 18.5 | yes |
| thbn_cp7 |  | 94 | 5.36±0.52 | 3.91±0.38 | 81 | 25.1 | 10400 | 0.274 | 26.4 | yes |
| thbn_cp8a | fraction 1 | 84 | 9.49±1.10 | 6.92±0.80 |  |  |  |  |  |  |
| thbn_cp8b | fraction 2 | 93 | 3.28±0.68 | 2.39±0.50 | 56 | 25.9 |  |  |  | yes |
| thbn_cp9a |  | 92.67 | 2.37±0.73 | 1.68±0.52 | 20 | 28.6 |  |  |  | yes |
| thbn_cp9b |  | 98.98 | 18.20±4.00 | 12.90±2.83 |  |  |  |  |  |  |
| thbn_cp10 |  | 90.8 | 1.78±0.68 | 1.27±0.48 | NA | >60 |  |  |  | yes |
| thbn_cp11 |  | 90 | 0.19±0.16 | 0.14±0.11 | 20 | 2.61 | 50600 | 0.135 | 2.7 | yes |
| thbn_cp12 |  | 92 | 0.91±0.17 | 0.66±0.12 | 20 | 3.5 |  |  |  | yes |
| thbn_cp13 |  | 86 | 11.03±2.76 | 8.04±2.02 |  |  |  |  |  |  |
| thbn_cp14 |  | 80 | 13.13±2.47 | 9.58±1.80 |  |  |  |  |  |  |
| thbn_cp15 |  | 99 | 0.23±0.14 | 0.17±0.10 | 20 | 1.3 | 115000 | 0.187 | 1.6 | yes |
| thbn_cp16 |  | 98 | 6.48±1.11 | 4.73±0.81 | 20 | 84.3 |  |  |  | yes |
| thbn_cp17 |  | 96.88 | 5.23±1.34 | 3.70±0.95 | 20 | >60 |  |  |  | yes |
| thbn_cp18 |  | 94 | 4.14±0.56 | 3.02±0.41 | 14 | 23.7 | 14400 | 0.348 | 24.1 | yes |
| thbn_cp19a |  | 52 | 0.17±0.14 | 0.13±0.11 |  |  |  |  |  |  |
| thbn_cp19b |  | 87 | 4.17±1.45 | 3.04±1.06 | 20 | 9.3 |  |  |  | yes |
| thbn_cp20 |  | 92.8 | 16.47±2.80 | 11.67±2.00 | 48 | 26.3 |  |  |  | yes |
| thbn_cp21 |  | 94 | NA | NA |  |  |  |  |  |  |
| thbn_cp22a |  | 92 | 6.23±2.64 | 4.55±1.93 | 20 | 10.9 |  |  |  | yes |
| thbn_cp22b |  | 99 | NA | NA |  |  |  |  |  |  |
| thbn_cp23 |  | 92 | NA | NA |  |  |  |  |  |  |

**Sup. Table 2.** MDM2:p53 inhibitors selected for re-synthesis and experimental characterization.

| *Name* | *fraction* | *LCMS Purity (%)* | *EC50 (μM)* | *NMR solubility (max. effective concentration μM)* | *steady-state KD (μM) (1mM TCEP)* | *steady-state KD (μM) (no TCEP)* | *Competitive with Nutlin-3A* |
| --- | --- | --- | --- | --- | --- | --- | --- |
| *Nutlin-3A* |  |  | 0.22±0.07 | 75 | 0.03 | 0.04 | yes |
| *M6 (ref)* |  | 99.08 | 32.01±7.49 | 20 | > 60 | 10.8 | yes |
| *mdm2_cp1* |  | 95.84 | NA |  |  |  |  |
| *mdm2_cp2* |  | 89.76 | 39.19±9.33 | 50 | 31.9 | 19.7 | yes |
| *mdm2_cp3* |  | 94.29 | NA |  |  |  |  |
| *mdm2_cp4a* | fraction 1 | 98.63 | 50.31±0.72 | 20 |  |  |  |
| *mdm2_cp4b* | fraction 2 | 92.69 | NA |  |  |  |  |
| *mdm2_cp5* |  | 99.75 | 5.73±4.01 | 20 | 3.4 | 2.1 | yes |
| *mdm2_cp6a* | fraction 1 | 91.28 | NA |  |  |  |  |
| *mdm2_cp6b* | fraction 2 | 91.59 | NA |  |  |  |  |
| *mdm2_cp7* |  | 97.49 | 44.58±6.35 | 50 | 90 | 135.0 | yes |
| *mdm2_cp8* |  | 93.3 | 6.87±1.27 | 50 | 18.9 | 19.1 | yes |
| *mdm2_cp9* |  | 99.8 | 2.4±1.51 | 20 | NA | 2.3 | yes |
| *mdm2_cp10* |  | 99.94 | NA |  |  |  |  |
| *mdm2_cp11a* | fraction 1 | 94.06 | 4.71±1.14 | 20 |  |  |  |
| *mdm2_cp11b* | fraction 2 | 92.92 | 3.79±1.01 | 20 | 1.4 | NA | NA |
| *mdm2_cp12* |  | 54.89 + 40.73 | 5.67±3.1 |  |  |  |  |
| *mdm2_cp13* |  | 99.69 | NA | 150 | 367.0 | 255.0 | yes |
| *mdm2_cp14* |  | 91.04 | 20.54±6.43 | 20 |  |  |  |
| *mdm2_cp15a* | fraction 1 | 99.68 | NA |  |  |  |  |
| *mdm2_cp15b* | fraction 2 | 97.12 | NA |  |  |  |  |
| *mdm2_cp16* |  | 96.03 | 25.52±0.26 | 20 | 36.4 | 11.5 | yes |
| *mdm2_cp17* |  | 83.61 + 98.78 | 30.85±1.93 | 150 |  |  |  |
| *mdm2_cp18a* | fraction 1 | 98.66 | NA |  |  |  |  |
| *mdm2_cp18b* | fraction 2 | 90.45 | NA |  |  |  |  |

## Supplementary Figures

**Sup. Fig. 1.**
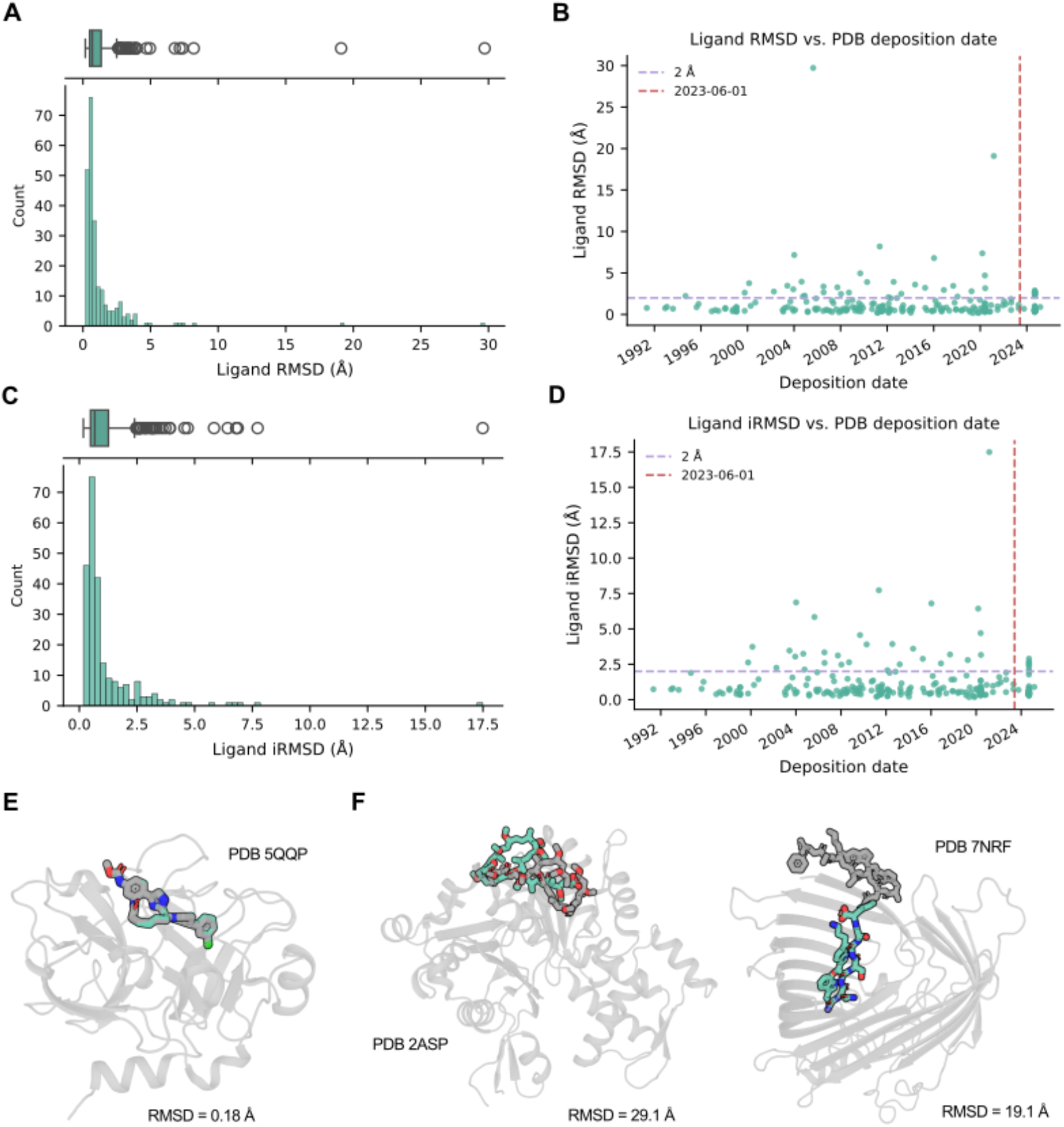
Boltz cofolding test on a macrocyclic ligand dataset **A:** Distribution of ligand RMSD values for Boltz cofolding on a benchmark set of 240 macrocycle-protein complexes^1^ (see Supplementary Methods). Most macrocycles are predicted accurately with ligand RMSD < 2 Å. **B:** Ligand RMSD versus PDB deposition date showing no clear correlation between deposition date and ligand RMSD. The dashed line indicates the training cutoff date. **C:** Distribution of ligand interface RMSD (iRMSD) values for the same benchmark set (see Supplementary Methods), indicating that the majority of predicted macrocycles correctly engage their target binding interface. **D:** Ligand interface RMSD (iRMSD) versus PDB deposition date, showing no correlation between deposition date and ligand interface RMSD. The dashed line indicates the training cutoff date. **E:** Example of a successfully predicted macrocycle pose (PDB 5QQP) with a near-native agreement between the predicted and experimental structures. **F:** Examples of failed predictions. Left: incorrect pose orientation (PDB 2ASP). Right: mislocalized binding pose (PDB 7NRF).

**Sup. Fig. 2.**
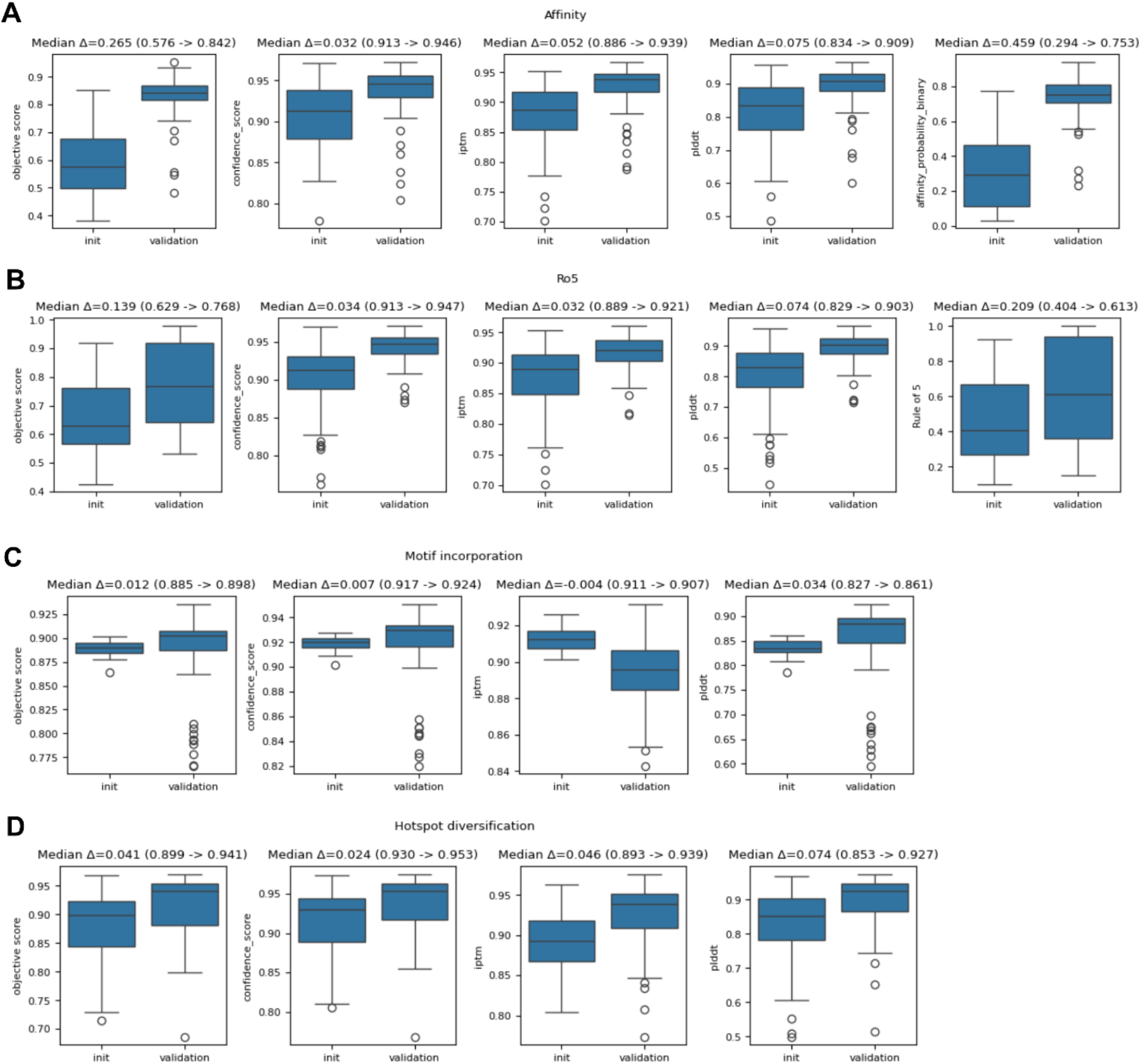
Score distributions for initial and final validation designs Distributions of co-optimized scores comparing randomly initialized designs (“init”) and final cyclic peptide ligands (“validation”). **A:** Co-optimization metrics, including confidence score, ipTM, mean ligand pLDDT, and binary probability affinity prediction evaluated over 100 independent trajectories with 100 optimization steps each. **B:** Co-optimization of beyond-Rule-of-5 (bRo5)–related metrics across 100 trajectories with 100 steps each. **C:** Motif incorporation scores across 50 trajectories with 100 steps each. **D:** Hotspot diversification score distributions across 50 trajectories with 100 steps each.

**Sup. Fig. 3.**
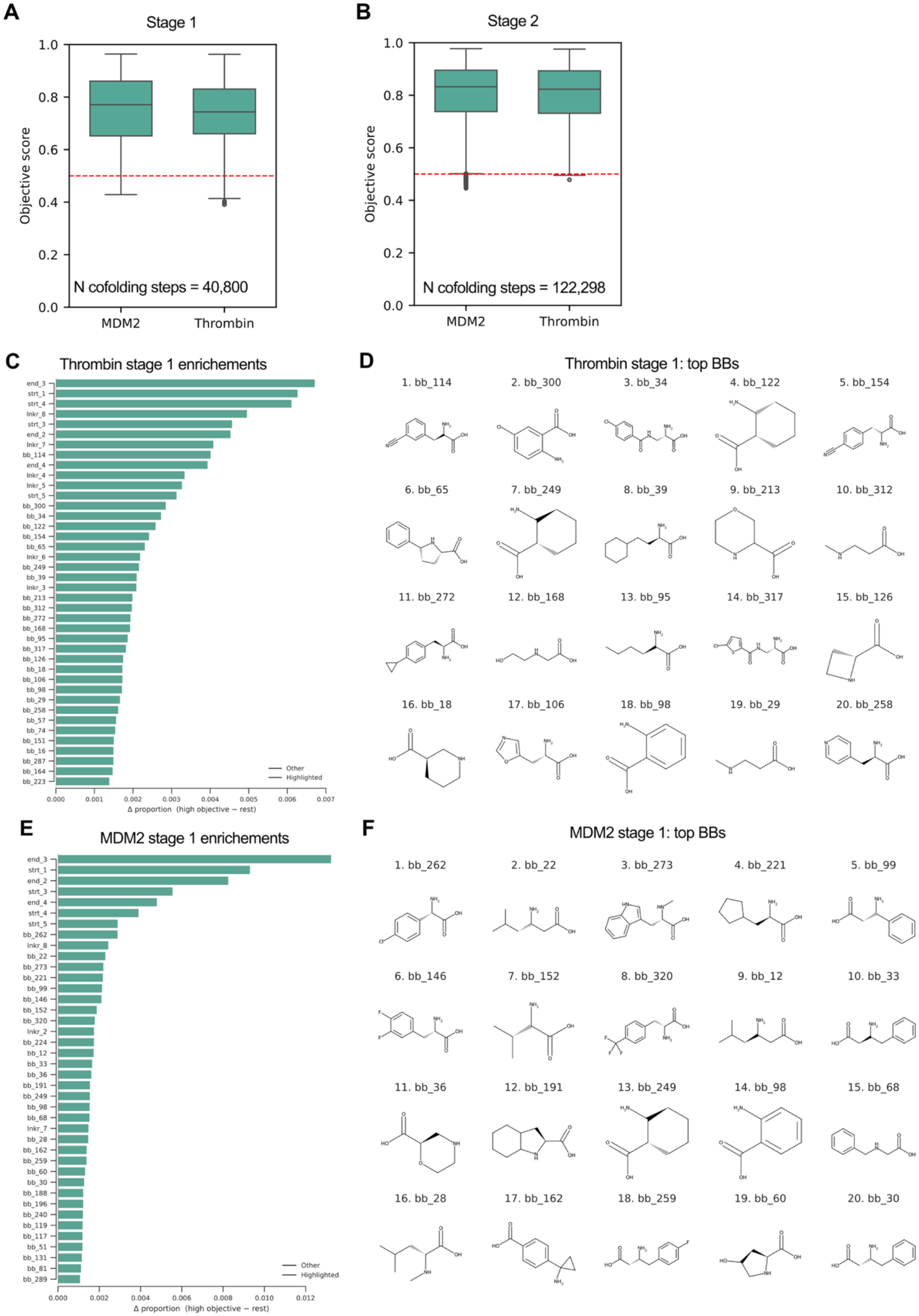
Stage 1 building block enrichments **A, B:** Distribution of objective scores for Stage 1 (A; full building block pool) and Stage 2 (B; reduced pool, 102 ncAAs). **C**: Top enriched building blocks for Thrombin in Stage 1. Enrichment (Δ proportion) is defined as the fraction of designs containing a given building block with objective score > 0.7 minus the fraction among designs with scores ≤ 0.7. Positive values indicate enrichment in high-scoring designs. **D**: Representative structures of building blocks enriched among high-scoring Thrombin designs. **E**: Top enriched building blocks for MDM2 in Stage 1. **F**: Representative structures of building blocks enriched among high-scoring MDM2 designs.

**Sup. Fig. 4.**
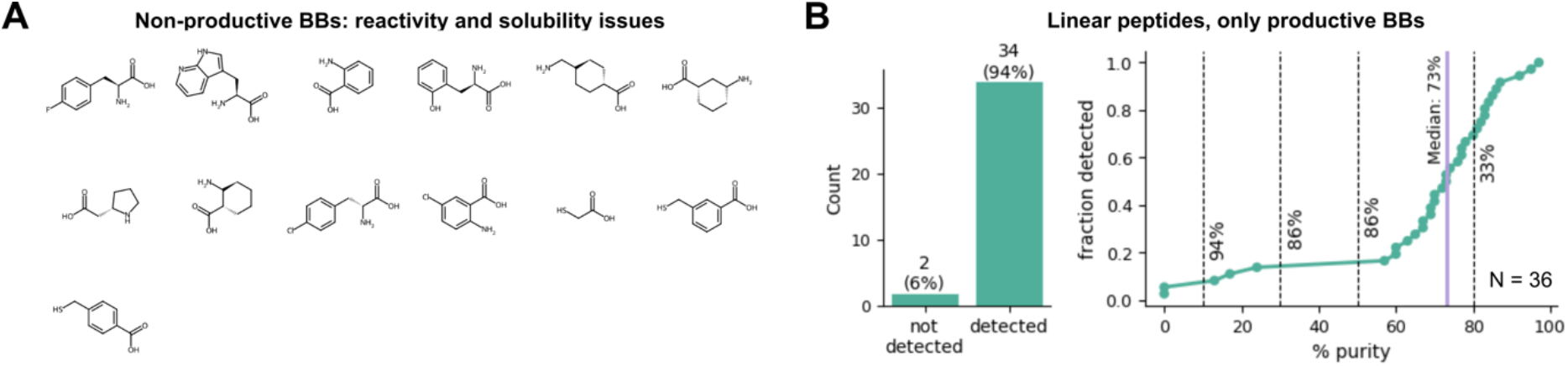
Non-productive building blocks and purity analysis of linear peptide library. **A:** Structures of non-productive building blocks associated with recurrent synthesis failures due to reactivity or solubility limitations. **B:** Linear peptides synthesized using only productive building blocks excluding building blocks with recurrent coupling or solubility issues, showing improved detection rates and higher median purity.

**Sup. Fig. 5.1.**
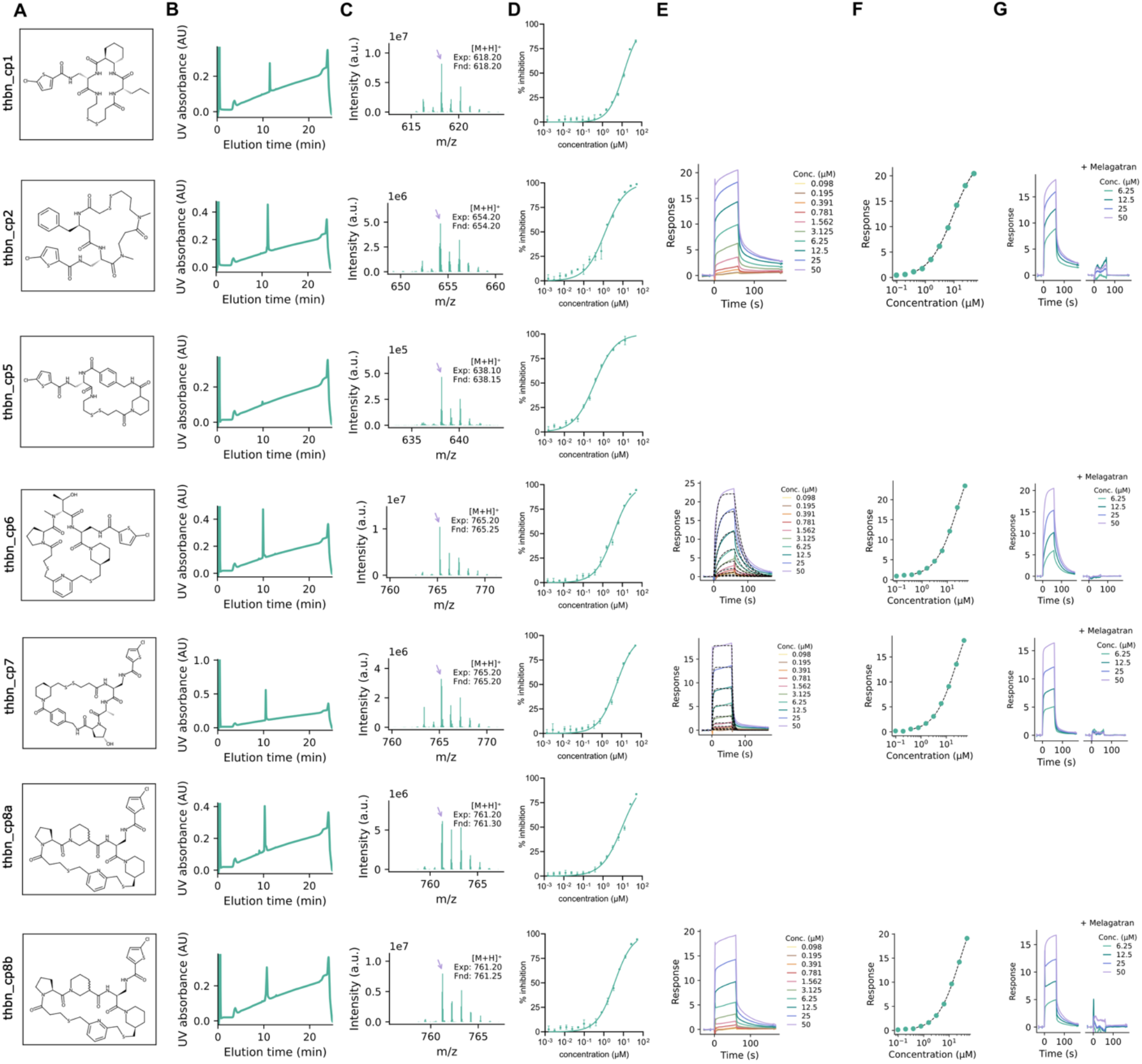
Chemical characterization and functional validation of selected Thrombin inhibitors **A:** Chemical structures of the designed cyclic peptide inhibitors. **B:** Analytical HPLC UV absorbance chromatograms (220 nm over 25 min) of purified compounds, confirming retention time and chromatographic purity. **C:** LC-MS spectra showing detection of the expected and found mass [M+H]^+^ for each compound. **D:** Dose-response curves from Thrombin inhibition assay, showing concentration-dependent inhibition. **E:** Representative SPR sensorgrams for selected compounds binding to immobilized Thrombin. **F:** Corresponding equilibrium binding fits derived from SPR measurements, yielding steady-state K_D_ values where possible. **G:** Competitive SPR sensorgrams performed in the absence and presence of an orthosteric Thrombin inhibitor Melagatran, assessing whether the designed cyclic peptides bind at the intended site on Thrombin.

**Sup. Fig. 5.2.**
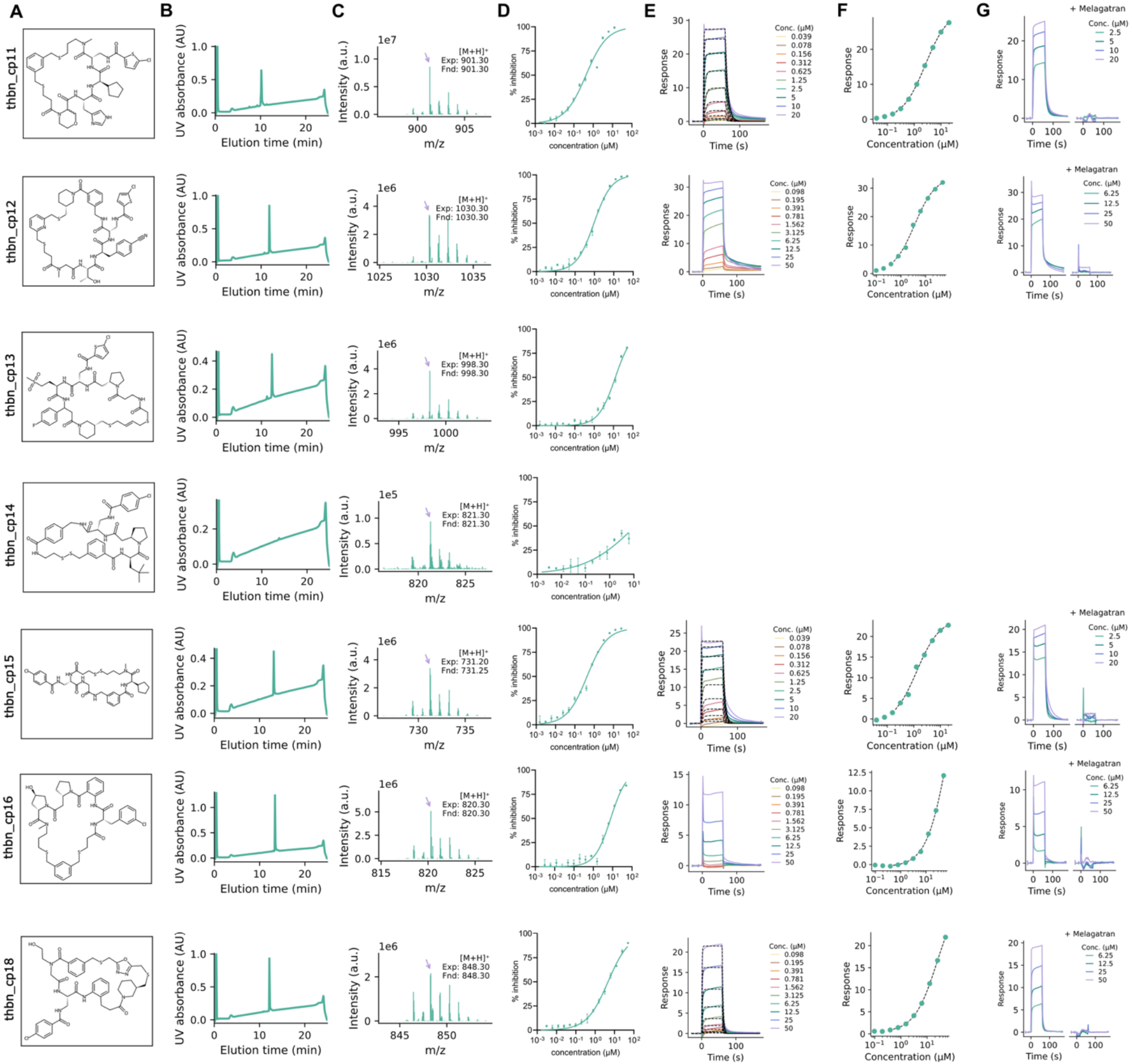
Chemical characterization and functional validation of selected Thrombin inhibitors **A:** Chemical structures of designed cyclic peptide inhibitors. **B:** Analytical HPLC UV absorbance chromatograms (220 nm over 25 min) of purified compounds, confirming retention time and chromatographic purity. **C:** LC-MS spectra showing detection of the expected and found mass [M+H]^+^ for each compound. **D:** Dose-response curves from Thrombin inhibition assay, showing concentration-dependent inhibition. **E:** Representative SPR sensorgrams for selected compounds binding to immobilized Thrombin. **F:** Corresponding equilibrium binding fits derived from SPR measurements, yielding steady-state K_D_ values. **G:** Competitive SPR sensorgrams performed in the absence and presence of an orthosteric Thrombin inhibitor Melagatran, assessing whether the designed cyclic peptides bind at the intended site on Thrombin.

**Sup. Fig. 5.3.**
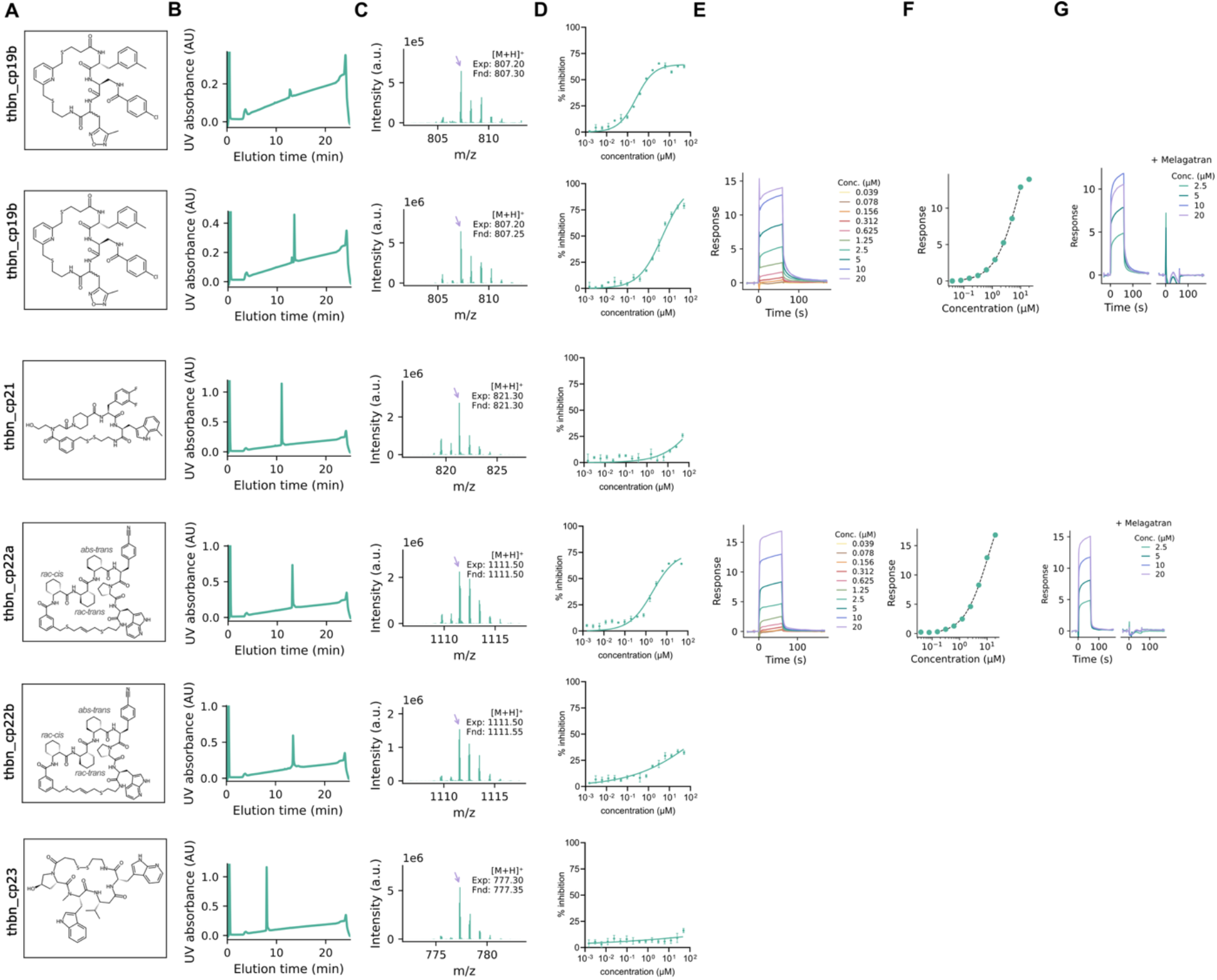
Chemical characterization and functional validation of selected Thrombin inhibitors **A:** Chemical structures of designed cyclic peptide inhibitors. **B:** Analytical HPLC UV absorbance chromatograms (220 nm over 25 min) of purified compounds, confirming retention time and chromatographic purity. **C:** LC-MS spectra showing detection of the expected and found mass [M+H]^+^ for each compound. **D:** Dose-response curves from Thrombin inhibition assay, showing concentration-dependent inhibition. **E:** Representative SPR sensorgrams for selected compounds binding to immobilized Thrombin. **F:** Corresponding equilibrium binding fits derived from SPR measurements, yielding steady-state K_D_ values. **G:** Competitive SPR sensorgrams performed in the absence and presence of an orthosteric Thrombin inhibitor Melagatran, assessing whether the designed cyclic peptides bind at the intended site on Thrombin.

**Sup. Fig. 5.4.**
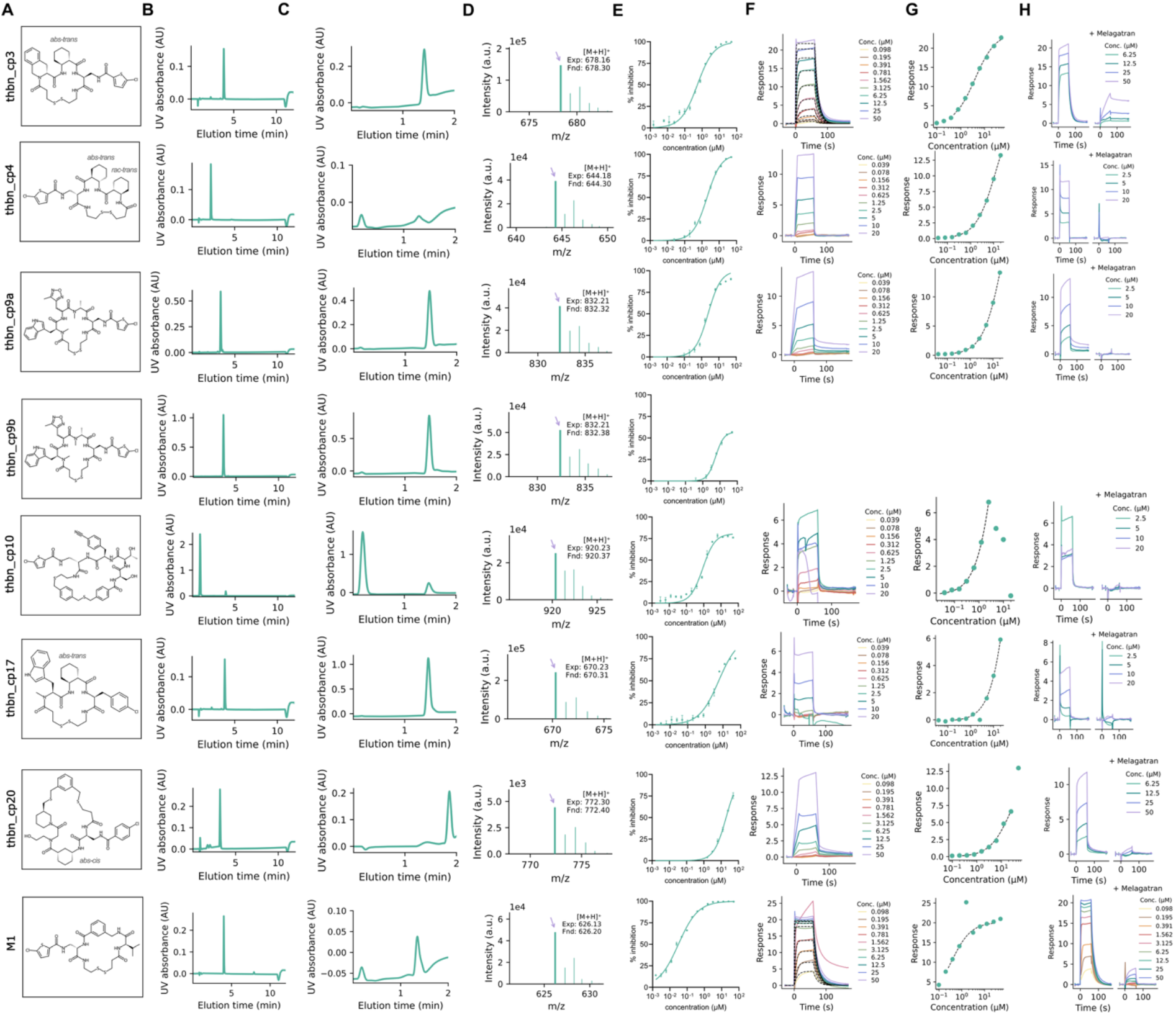
Chemical characterization and functional validation of selected Thrombin inhibitors **A:** Chemical structures of purified cyclic peptide inhibitors. The M1 compound is a positive control from Habeshian et al.^3^ **B:** Analytical HPLC UV absorbance chromatograms (220 nm over 10 min) of purified compounds, confirming retention time and chromatographic purity. **C:** LC-MS UV chromatograms (220 nm over 2 min) for molecular weight confirmation. **D:** LC-MS spectra showing detection of the expected and found mass [M+H]^+^ for each compound. **E:** Dose-response curves from the Thrombin inhibition assay, showing concentration-dependent inhibition. **F:** SPR sensorgrams for selected compounds binding to immobilized Thrombin. **G:** Corresponding equilibrium binding fits derived from SPR measurements, yielding steady-state K_D_ values. **H:** Competitive SPR measurements performed in the absence and presence of an orthosteric Thrombin inhibitor Melagatran, assessing whether the designed cyclic peptides bind at the intended site on Thrombin.

**Sup. Fig. 6.1.**
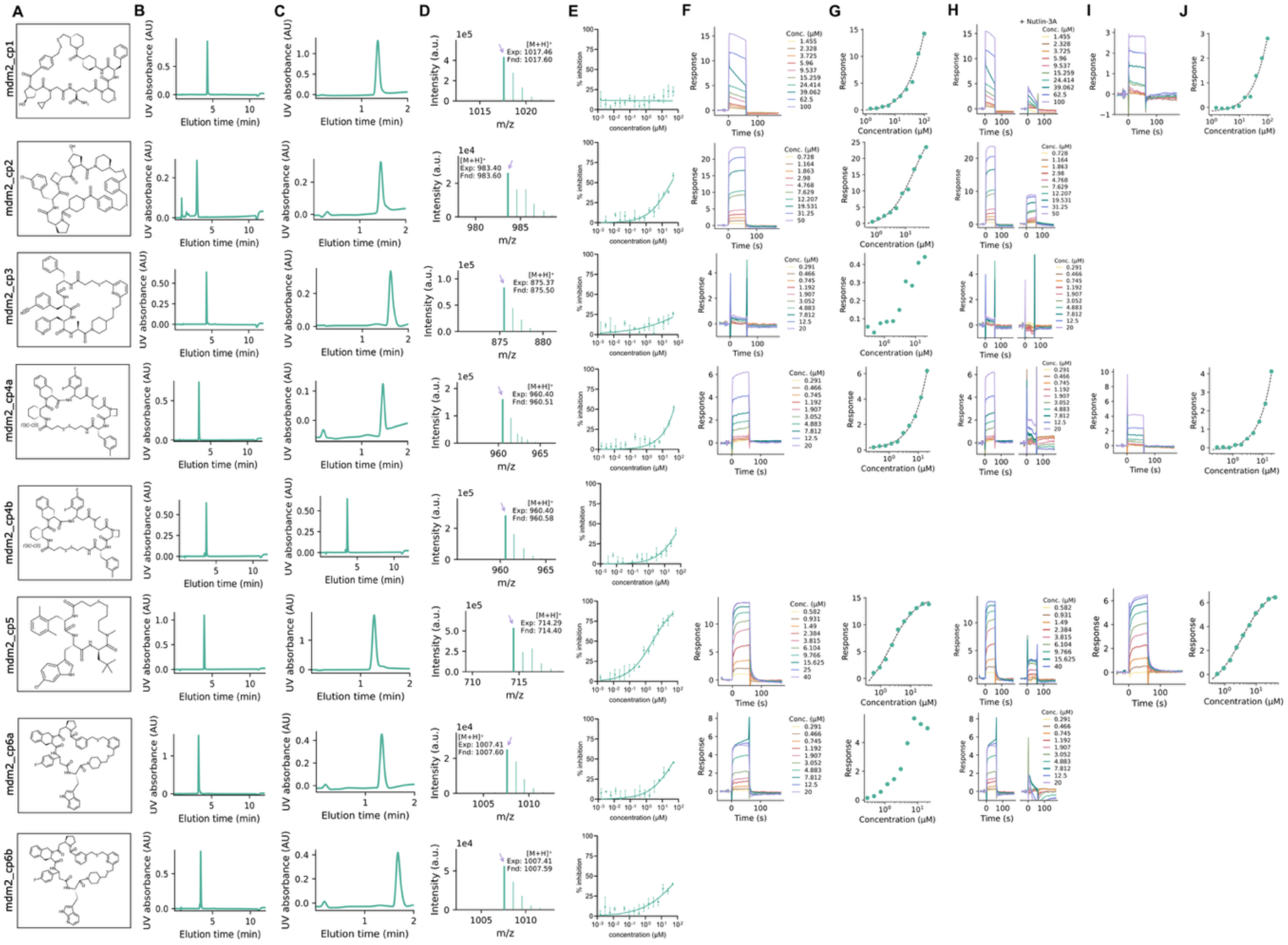
Chemical characterization and functional validation of selected MDM2:p53 inhibitors **A:** Chemical structures of purified cyclic peptide inhibitors. **B:** Analytical HPLC UV absorbance chromatograms (220 nm over 10 min) of purified compounds, confirming retention time and chromatographic purity. **C:** LC-MS UV chromatograms (220 nm over 2 min) for molecular weight confirmation. **D:** LC-MS spectra showing detection of the expected and found mass [M+H]^+^ for each compound. **E:** Dose-response curves from the MDM2:p53 HTRF competition assay, showing concentration-dependent disruption of the MDM2:p53 interaction. **F:** SPR sensorgrams for selected compounds binding to immobilized MDM2. **G:** Corresponding equilibrium binding fits derived from SPR measurements, yielding steady-state K_D_ values. **H:** Competitive SPR measurements performed in the absence and presence of an orthosteric MDM2 inhibitor Nutlin-3A, assessing whether the designed cyclic peptides bind at the intended site on MDM2. **I:** SPR sensorgrams for disulfide-cyclized peptides measured under reducing conditions using TCEP, which is expected to yield the corresponding linear peptide forms. **J:** Corresponding equilibrium binding fits from SPR measurements under reducing conditions, showing the concentration-dependent binding response of the expected linear peptides.

**Sup. Fig. 6.2.**
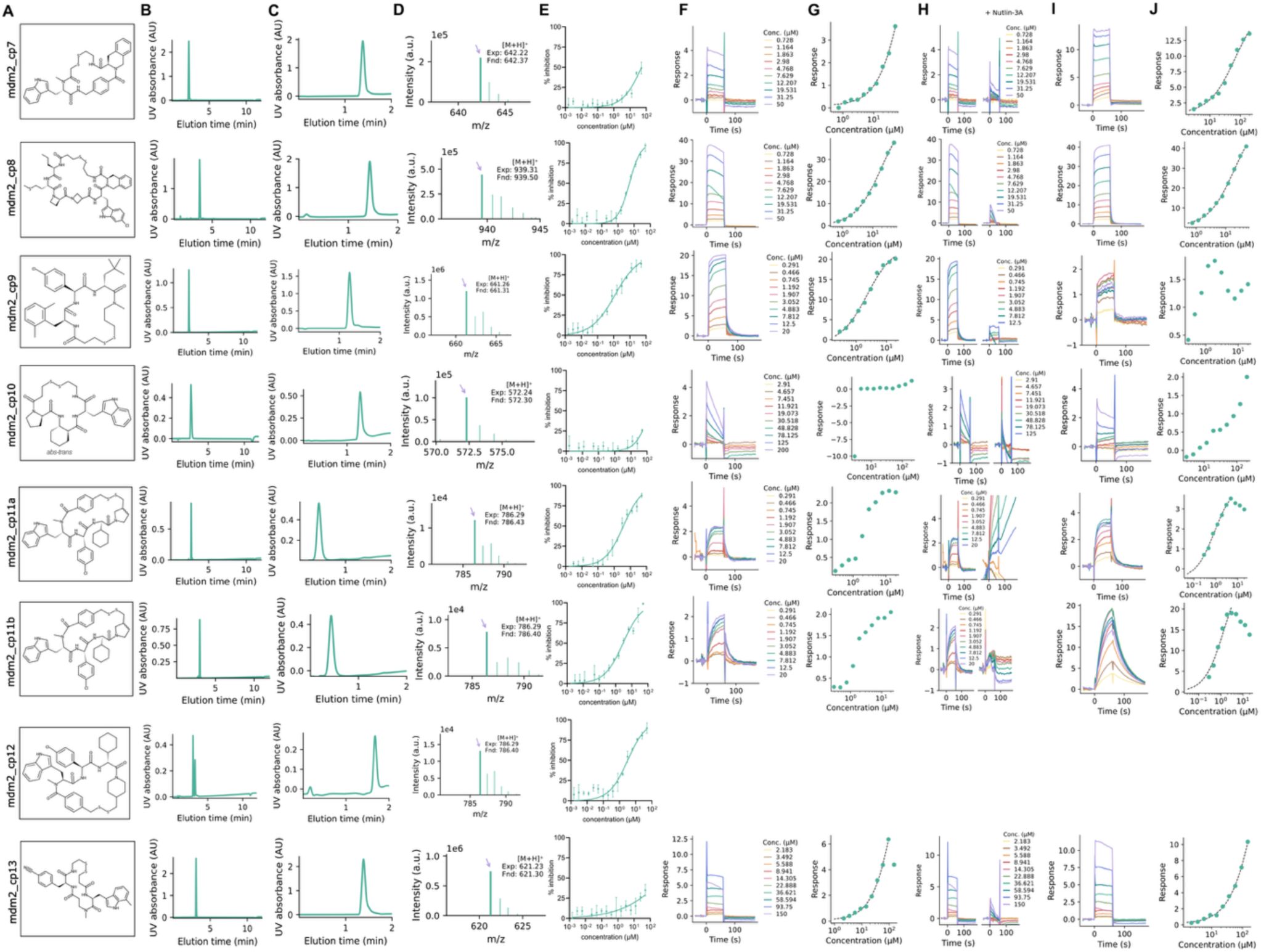
Chemical characterization and functional validation of selected MDM2:p53 inhibitors **A:** Chemical structures of purified cyclic peptide inhibitors. **B:** Analytical HPLC UV absorbance chromatograms (220 nm over 10 min) of purified compounds, confirming retention time and chromatographic purity. **C:** LC-MS UV chromatograms (220 nm over 2 min) for molecular weight confirmation. **D:** LC-MS spectra showing detection of the expected and found mass [M+H]^+^ for each compound. **E:** Dose-response curves from the MDM2:p53 HTRF competition assay, showing concentration-dependent disruption of the MDM2:p53 interaction. **F:** SPR sensorgrams for selected compounds binding to immobilized MDM2. **G:** Corresponding equilibrium binding fits derived from SPR measurements, yielding steady-state K_D_ values. **H:** Competitive SPR measurements performed in the absence and presence of an orthosteric MDM2 inhibitor Nutlin-3A, assessing whether the designed cyclic peptides bind at the intended site on MDM2. **I:** SPR sensorgrams for disulfide-cyclized peptides measured under reducing conditions using TCEP, which is expected to yield the corresponding linear peptide forms. **J:** Corresponding equilibrium binding fits from SPR measurements under reducing conditions, showing the concentration-dependent binding response of the expected linear peptides.

**Sup. Fig. 6.3.**
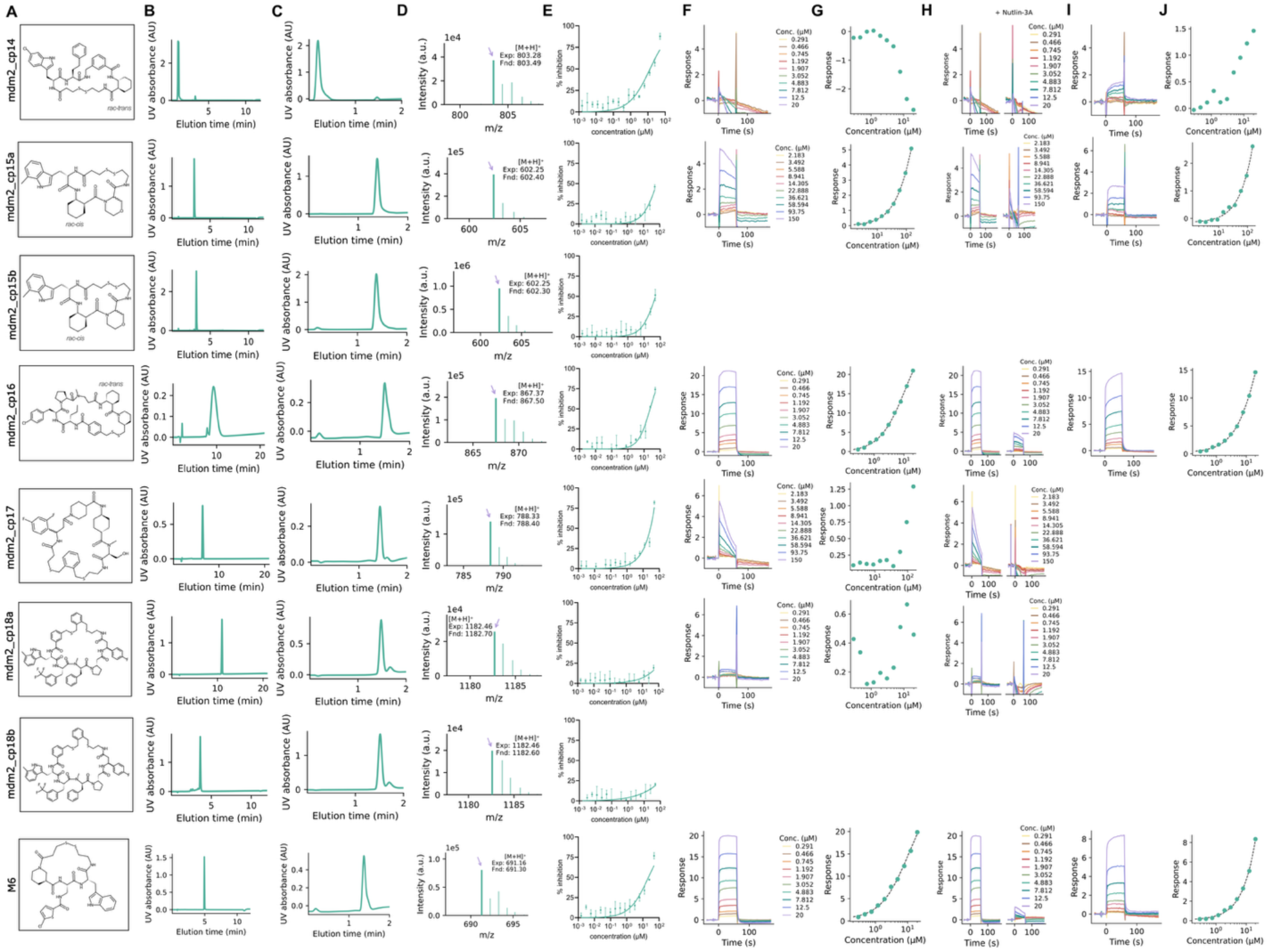
Chemical characterization and functional validation of selected MDM2:p53 inhibitors **A:** Chemical structures of purified cyclic peptide inhibitors. The M6 compound is a positive control from Habeshian et al.^3^ **B:** Analytical HPLC UV absorbance chromatograms (220 nm over 10 min) of purified compounds, confirming retention time and chromatographic purity. **C:** LC-MS UV chromatograms (220 nm over 2 min) for molecular weight confirmation. **D:** LC-MS spectra showing detection of the expected and found mass [M+H]^+^ for each compound. **E:** Dose-response curves from the MDM2:p53 HTRF competition assay, showing concentration-dependent disruption of the MDM2:p53 interaction. **F:** SPR sensorgrams for selected compounds binding to immobilized MDM2. **G:** Corresponding equilibrium binding fits derived from SPR measurements, yielding steady-state K_D_ values. **H:** Competitive SPR measurements performed in the absence and presence of an orthosteric MDM2 inhibitor Nutlin-3A, assessing whether the designed cyclic peptides bind at the intended site on MDM2. **I:** SPR sensorgrams for disulfide-cyclized peptides measured under reducing conditions using TCEP, which is expected to yield the corresponding linear peptide forms. **J:** Corresponding equilibrium binding fits from SPR measurements under reducing conditions, showing the concentration-dependent binding response of the expected linear peptide.

**Sup. Fig. 7.**
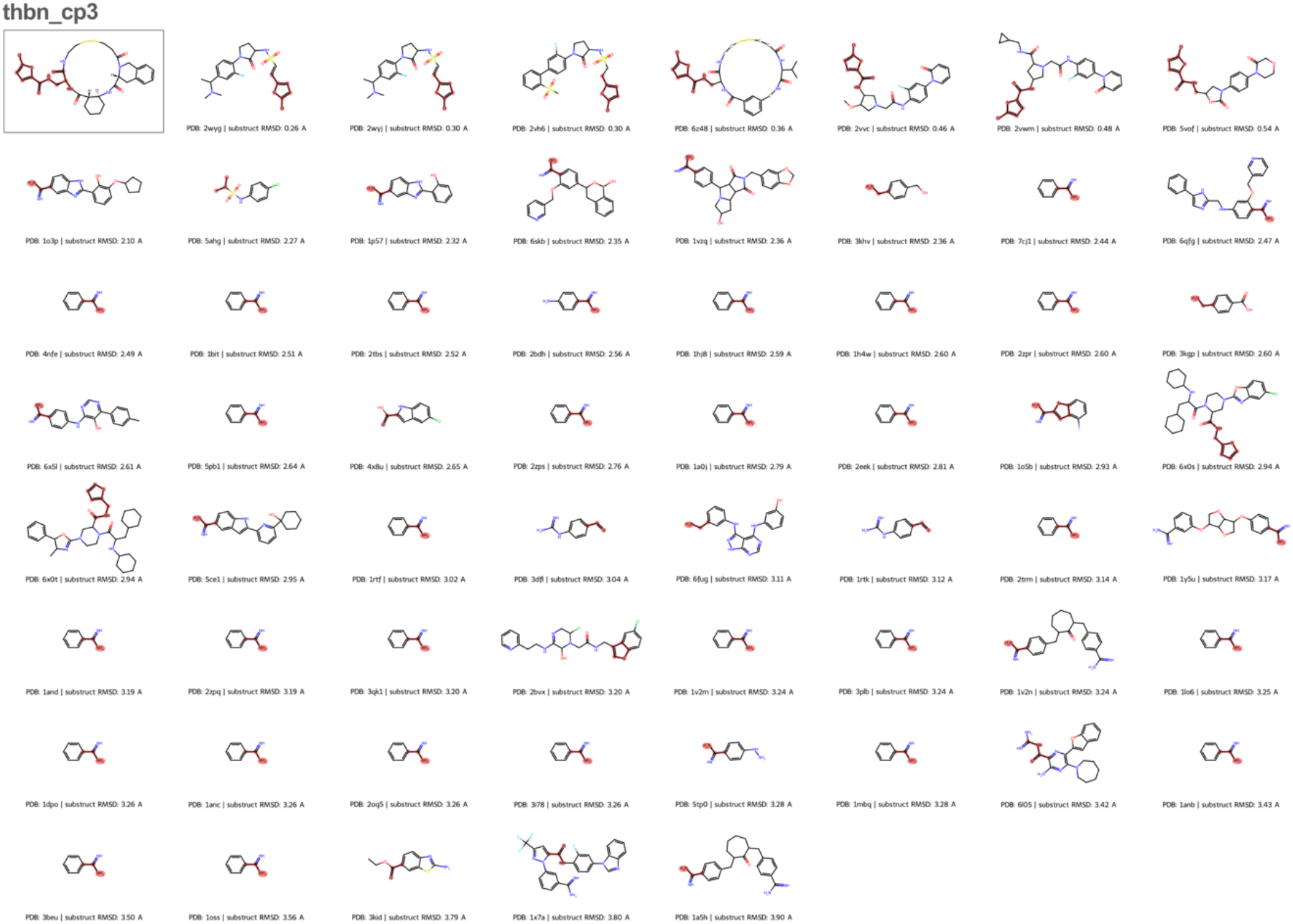
Substructure matches of the chlorothiophene moiety in thbn_cp3 The top-left boxed structure shows the designed cyclic peptide, with the query substructure highlighted. The remaining structures are Thrombin-bound PDB ligands that share a substructure with the query, ordered by increasing RMSD of the building block MCS. Highlighted atoms denote the MCS shared between the building block and each reference ligand.

**Sup. Fig. 8.**
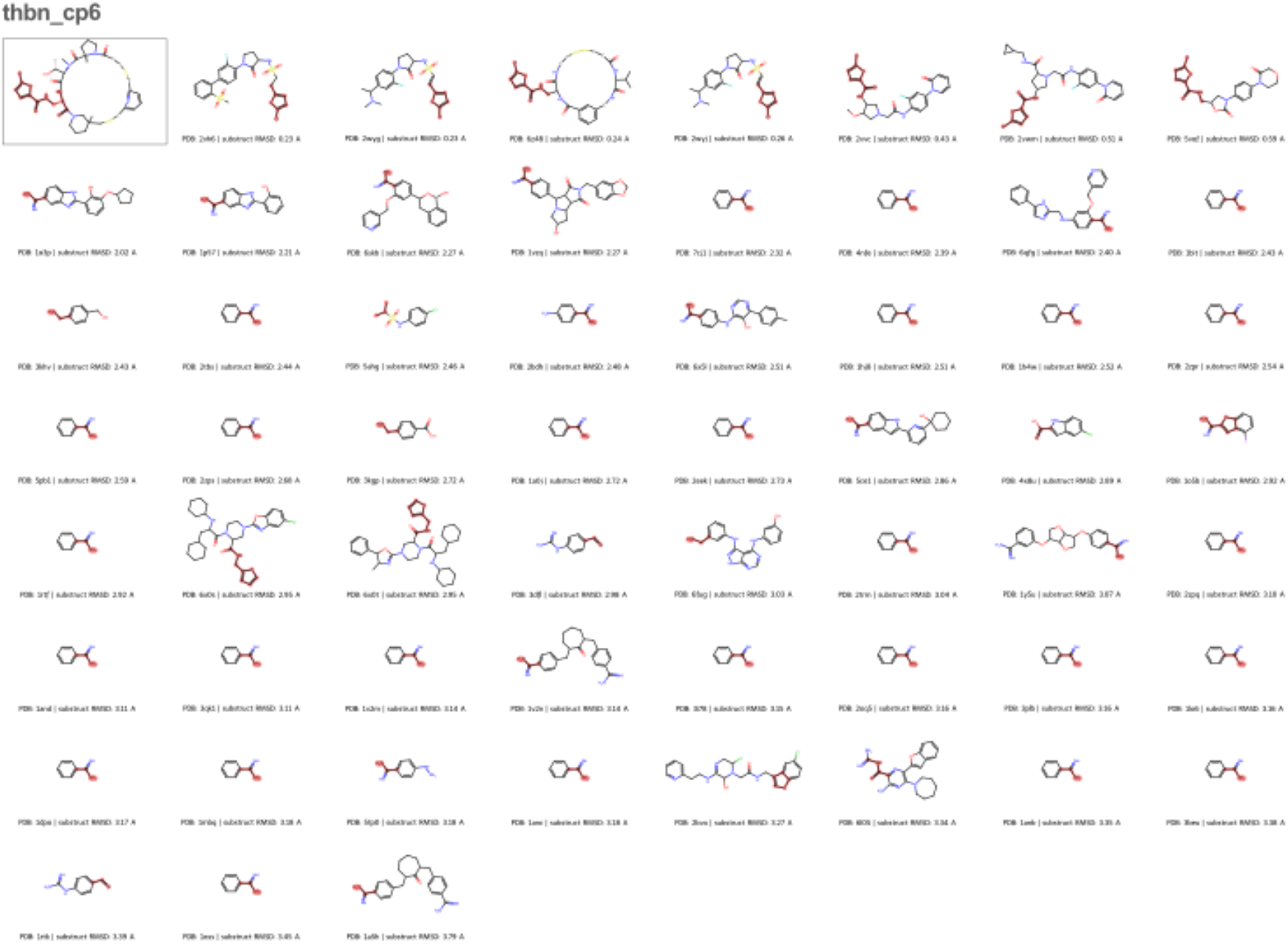
Substructure matches of the chlorothiophene moiety in thbn_cp6 The top-left boxed structure shows the designed cyclic peptide, with the query substructure highlighted. The remaining structures are Thrombin-bound PDB ligands that share a substructure with the query, ordered by increasing RMSD of the building block MCS. Highlighted atoms denote the MCS shared between the building block and each reference ligand.

**Sup. Fig. 9.**
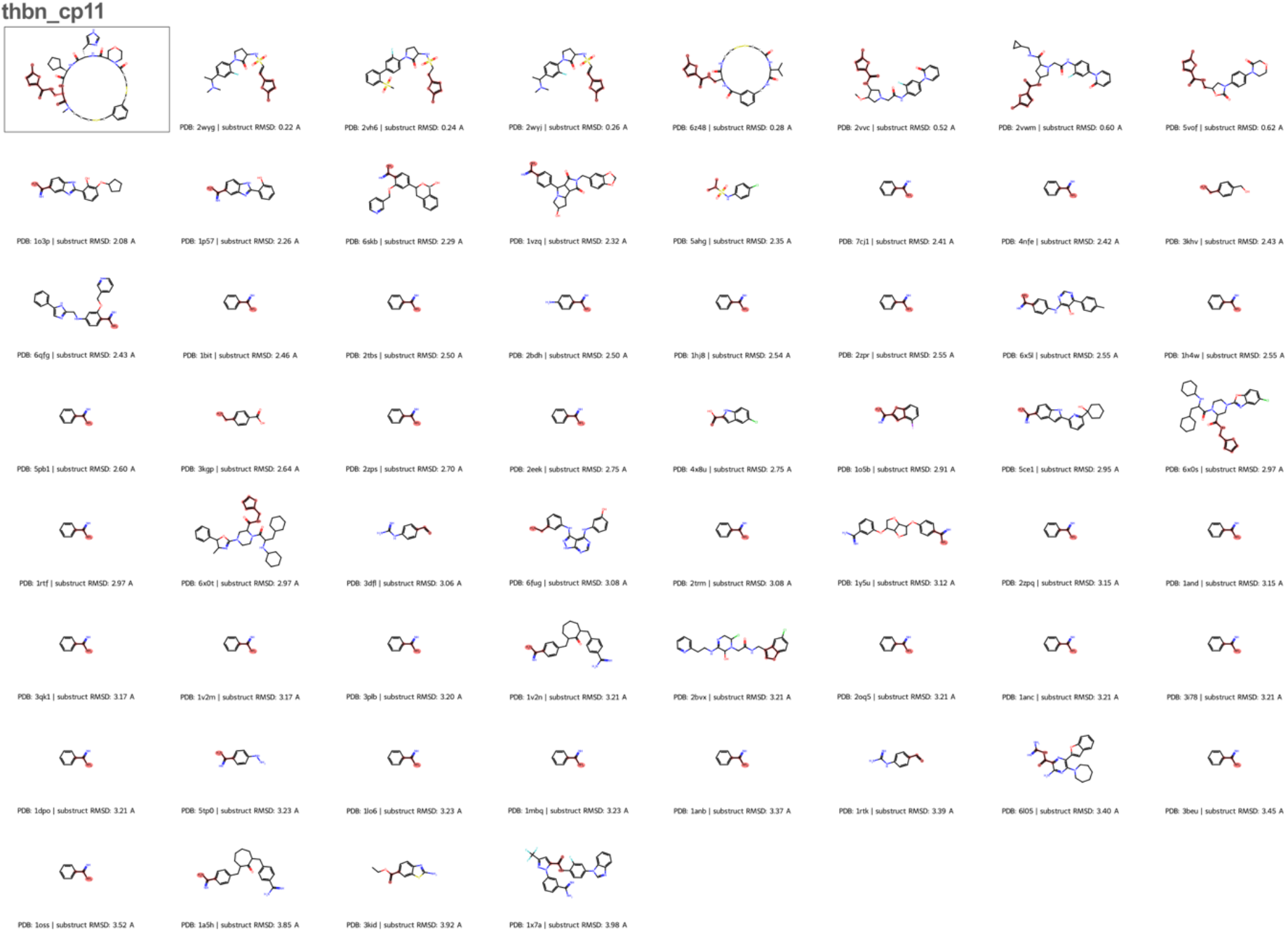
Substructure matches of the chlorothiophene moiety in thbn_cp11 The top-left boxed structure shows the designed cyclic peptide, with the query substructure highlighted. The remaining structures are Thrombin-bound PDB ligands that share a substructure with the query, ordered by increasing RMSD of the building block MCS. Highlighted atoms denote the MCS shared between the building block and each reference ligand.

**Sup. Fig. 10.**
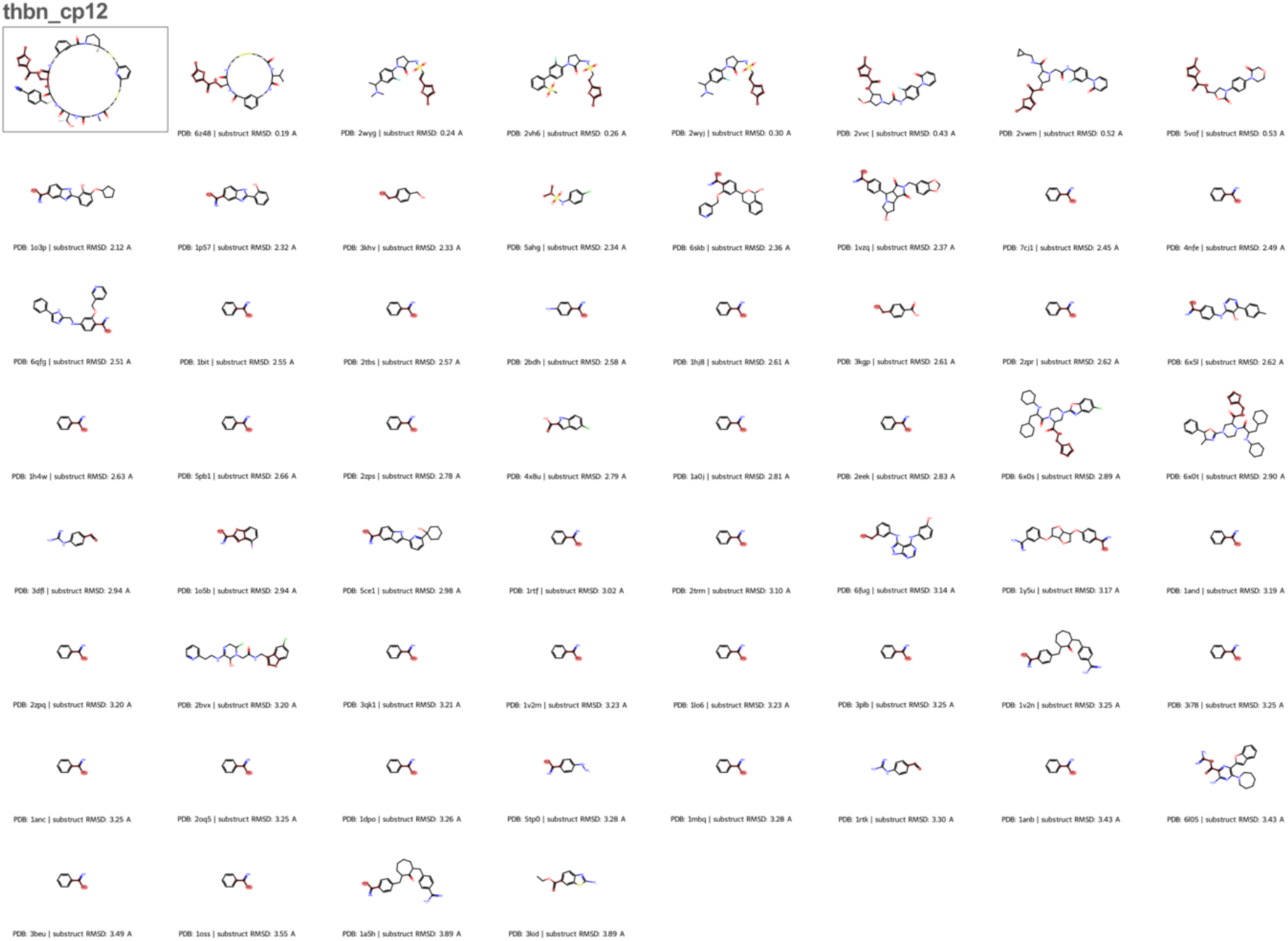
Substructure matches of the chlorothiophene moiety in thbn_cp12 The top-left boxed structure shows the designed cyclic peptide, with the query substructure highlighted. The remaining structures are Thrombin-bound PDB ligands that share a substructure with the query, ordered by increasing RMSD of the building block MCS. Highlighted atoms denote the MCS shared between the building block and each reference ligand.

**Sup. Fig. 11.**
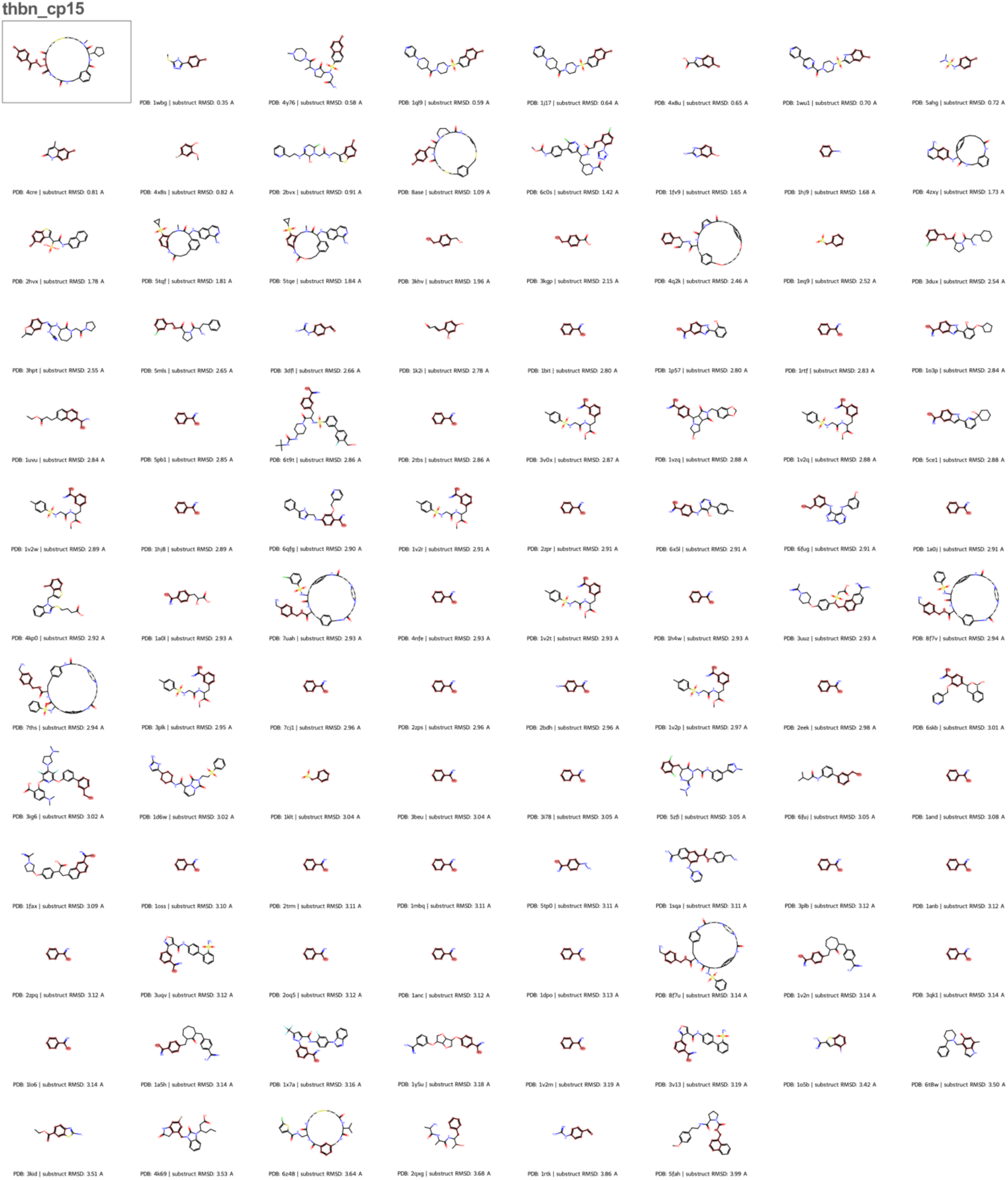
Substructure matches of the chlorophenyl moiety in thbn_cp15 The top-left boxed structure shows the designed cyclic peptide, with the query substructure highlighted. The remaining structures are Thrombin-bound PDB ligands that share a substructure with the query, ordered by increasing RMSD of the building block MCS. Highlighted atoms denote the MCS shared between the building block and each reference ligand.

**Sup. Fig. 12.**
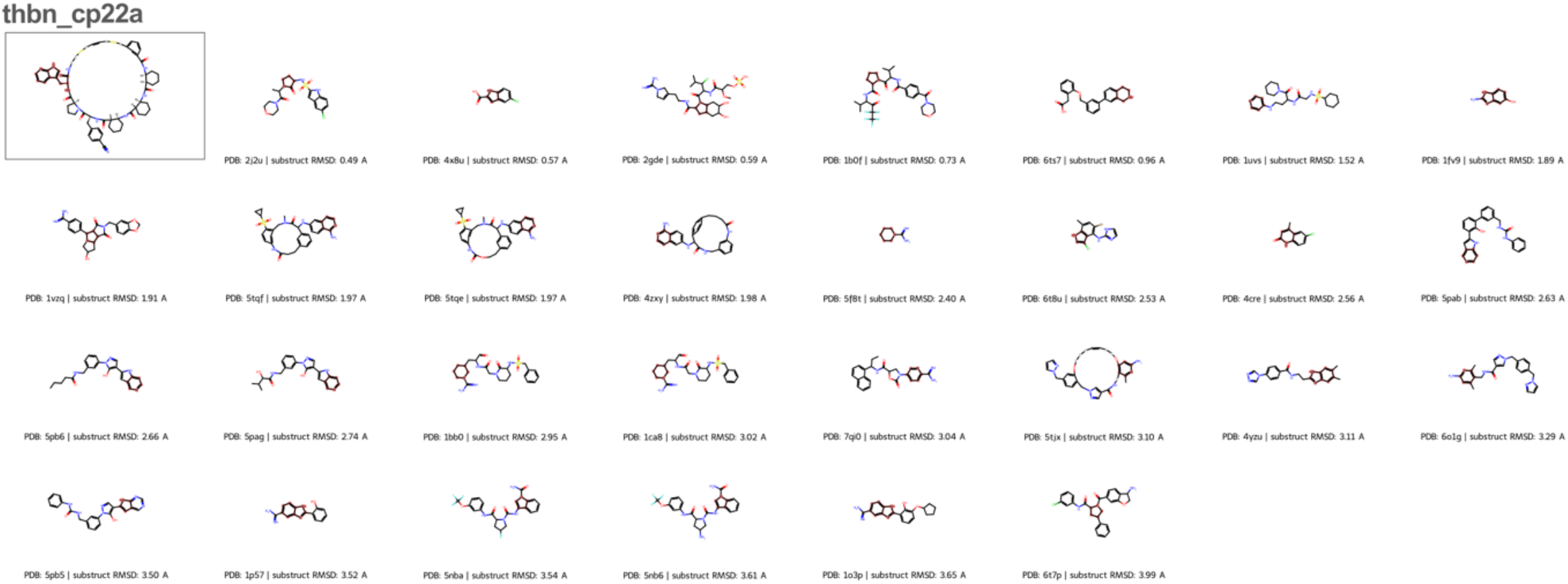
Substructure matches of the azatryptophan moiety in thbn_cp22a The top-left boxed structure shows the designed cyclic peptide, with the query substructure highlighted. The remaining structures are Thrombin-bound PDB ligands that share a substructure with the query, ordered by increasing RMSD of the building block MCS. Highlighted atoms denote the MCS shared between the building block and each reference ligand.

**Sup. Fig. 13.**
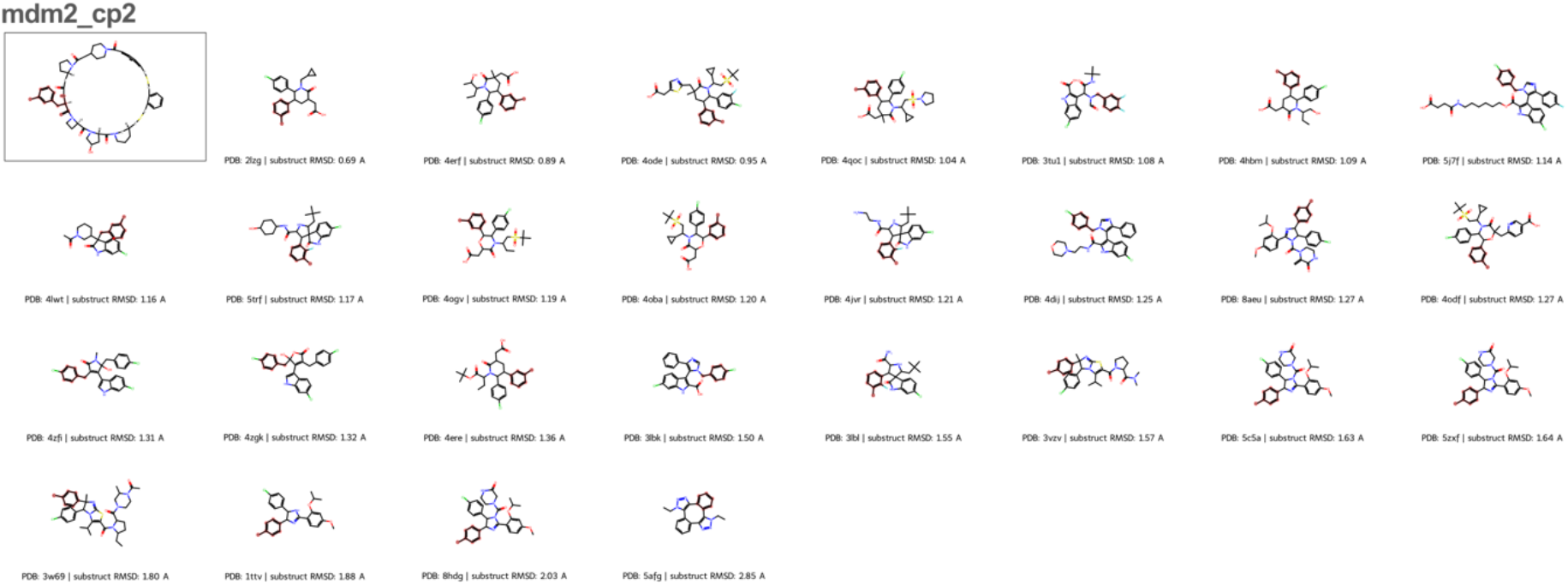
Substructure matches of the chlorophenyl moiety in mdm2_cp2 The top-left boxed structure shows the designed cyclic peptide, with the query substructure highlighted. The remaining structures are MDM2-bound PDB ligands that share a substructure with the query, ordered by increasing RMSD of the building block MCS. Highlighted atoms denote the MCS shared between the building block and each reference ligand.

**Sup. Fig. 14.**
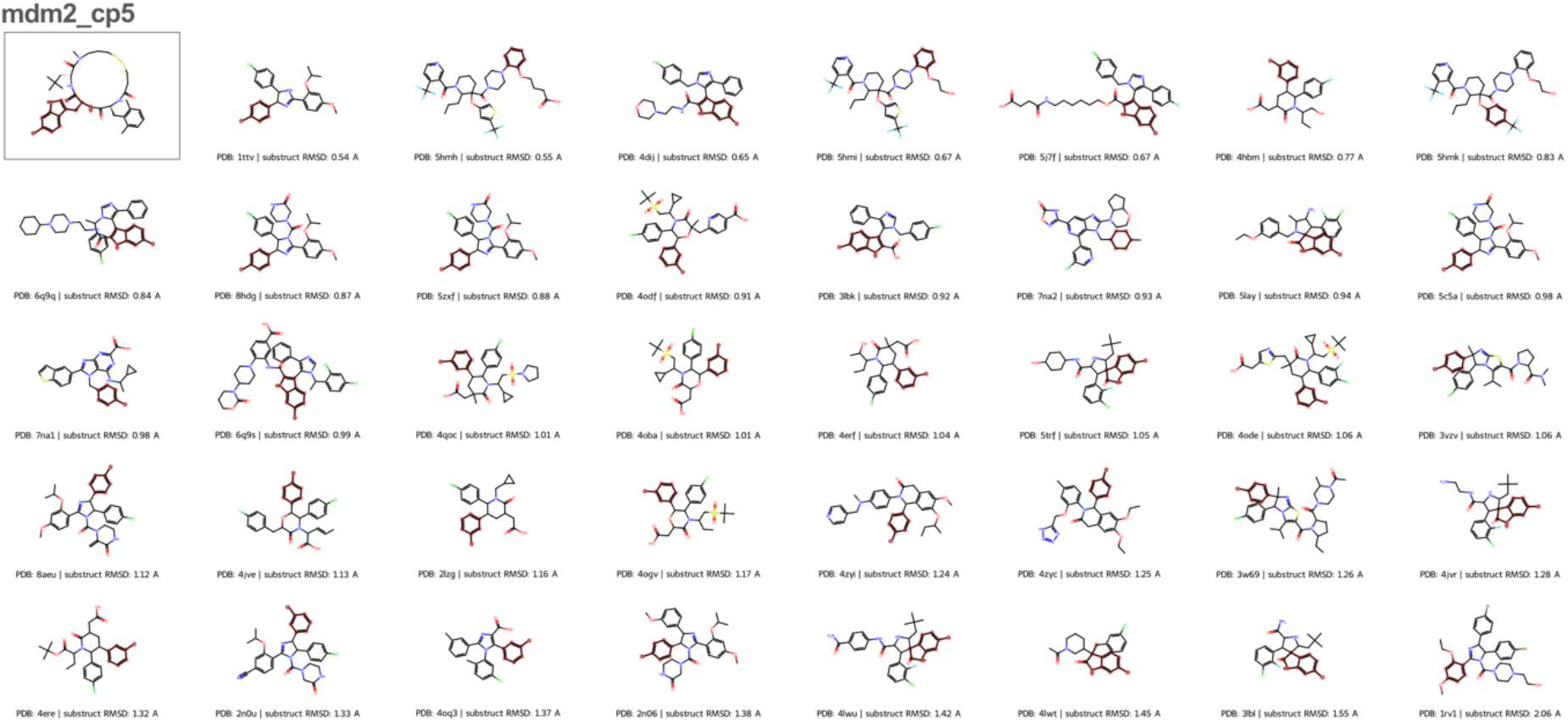
Substructure matches of the chlorotryptophan moiety in mdm2_cp5 The top-left boxed structure shows the designed cyclic peptide, with the query substructure highlighted. The remaining structures are MDM2-bound PDB ligands that share a substructure with the query, ordered by increasing RMSD of the building block MCS. Highlighted atoms denote the MCS shared between the building block and each reference ligand.

**Sup. Fig. 15.**
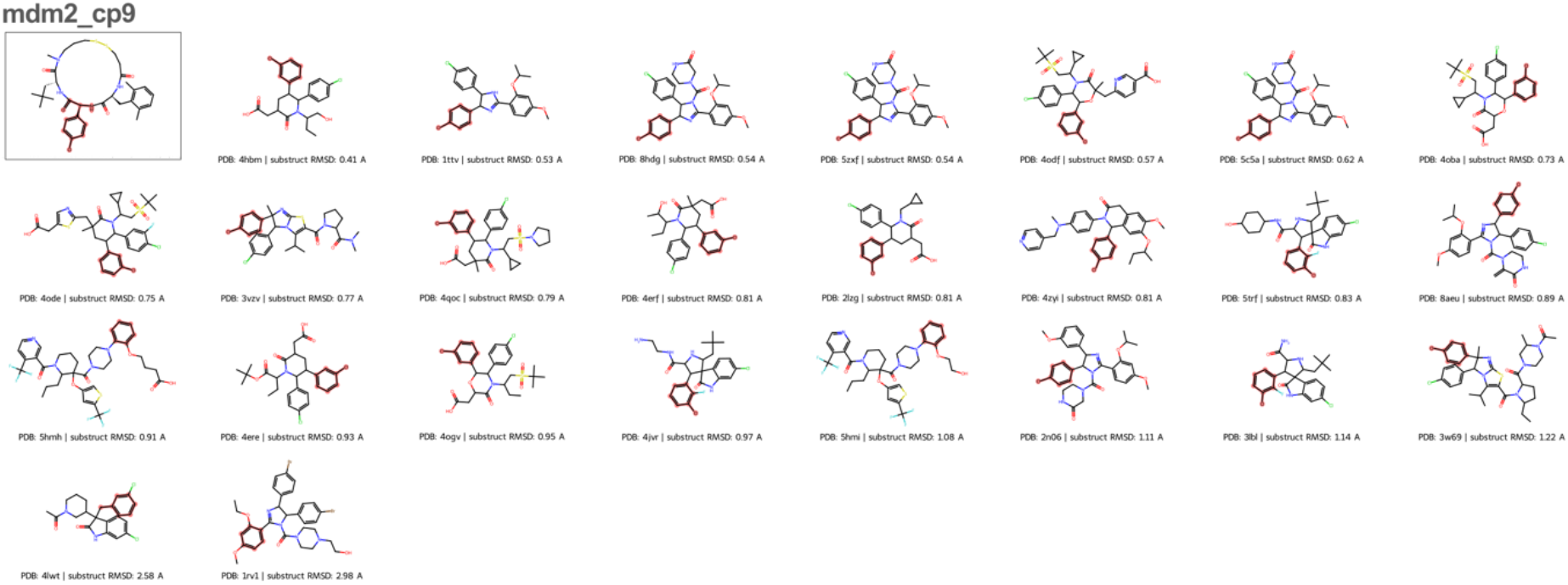
Substructure matches of the chlorophenyl moiety in mdm2_cpG The top-left boxed structure shows the designed cyclic peptide, with the query substructure highlighted. The remaining structures are MDM2-bound PDB ligands that share a substructure with the query, ordered by increasing RMSD of the building block MCS. Highlighted atoms denote the MCS shared between the building block and each reference ligand.

**Sup. Fig. 16.**
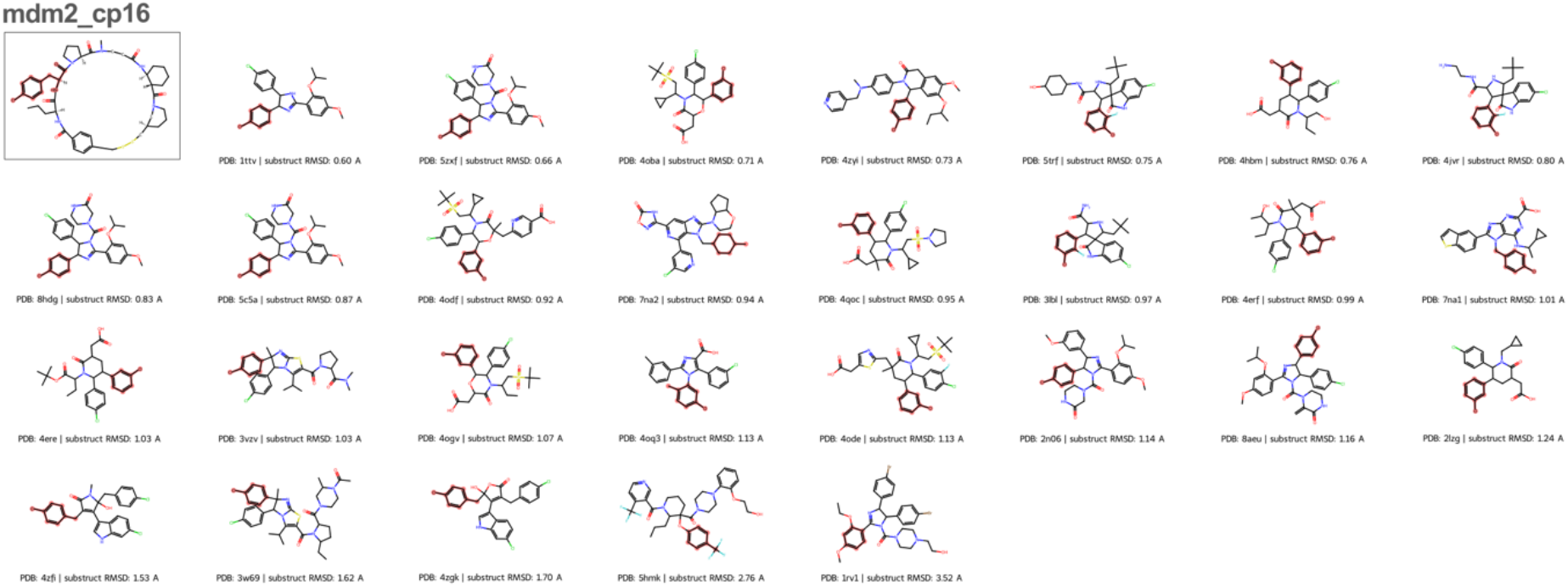
Substructure matches of the chlorophenyl moiety in mdm2_cp16 The top-left boxed structure shows the designed cyclic peptide, with the query substructure highlighted. The remaining structures are MDM2-bound PDB ligands that share a substructure with the query, ordered by increasing RMSD of the building block MCS. Highlighted atoms denote the MCS shared between the building block and each reference ligand.

## References

1. Hopkins, A. L. C Groom, C. R. The druggable genome. Nat. Rev. Drug Discov. 1, 727– 730 (2002).

2. Scott, D. E., Bayly, A. R., Abell, C. C Skidmore, J. Small molecules, big targets: drug discovery faces the protein–protein interaction challenge. Nat. Rev. Drug Discov. 15, 533–550 (2016).

3. Wells, J. A. C McClendon, C. L. Reaching for high-hanging fruit in drug discovery at protein–protein interfaces. Nature 450, 1001–1009 (2007).

4. Huggins, D. J., Sherman, W. C Tidor, B. Rational Approaches to Improving Selectivity in Drug Design. J. Med. Chem. 55, 1424–1444 (2012).

5. Anselmo, A. C., Gokarn, Y. C Mitragotri, S. Non-invasive delivery strategies for biologics. Nat. Rev. Drug Discov. 18, 19–40 (2019).

6. Muttenthaler, M., King, G. F., Adams, D. J. C Alewood, P. F. Trends in peptide drug discovery. Nat. Rev. Drug Discov. 20, 309–325 (2021).

7. Villar, E. A. et al. How proteins bind macrocycles. Nat. Chem. Biol. 10, 723–731 (2014).

8. Dougherty, P. G., Qian, Z. C Pei, D. Macrocycles as protein–protein interaction inhibitors. Biochem. J. 474, 1109–1125 (2017).

9. Ji, X., Nielsen, A. L. C Heinis, C. Cyclic Peptides for Drug Development. Angew. Chem. Int. Ed. 63, e202308251 (2024).

10. Kingwell, K. Macrocycle drugs serve up new opportunities. Nat. Rev. Drug Discov. 22, 771–773 (2023).

11. Deyle, K., Kong, X.-D. C Heinis, C. Phage Selection of Cyclic Peptides for Application in Research and Drug Development. Acc. Chem. Res. 50, 1866–1874 (2017).

12. Passioura, T., Liu, W., Dunkelmann, D., Higuchi, T. C Suga, H. Display Selection of Exotic Macrocyclic Peptides Expressed under a Radically Reprogrammed 23 Amino Acid Genetic Code. J. Am. Chem. Soc. 140, 11551–11555 (2018).

13. Habeshian, S. et al. Synthesis and direct assay of large macrocycle diversities by combinatorial late-stage modification at picomole scale. Nat. Commun. 13, 3823 (2022).

14. Merz, M. L. et al. De novo development of small cyclic peptides that are orally bioavailable. Nat. Chem. Biol. 20, 624–633 (2024).

15. Ji, X. et al. Generation of membrane-permeable cyclic peptides inhibiting protein– protein interaction. Nat. Chem. Biol. 1–11 (2026) doi:10.1038/s41589-026-02237-7.

16. Vinogradov, A. A., Yin, Y. C Suga, H. Macrocyclic Peptides as Drug Candidates: Recent Progress and Remaining Challenges. J. Am. Chem. Soc. 141, 4167–4181 (2019).

17. Hickey, J. L., Sindhikara, D., Zultanski, S. L. C Schultz, D. M. Beyond 20 in the 21st Century: Prospects and Challenges of Non-canonical Amino Acids in Peptide Drug Discovery. ACS Med. Chem. Lett. 14, 557–565 (2023).

18. Wilcken, R., Zimmermann, M. O., Lange, A., Joerger, A. C. C Boeckler, F. M. Principles and Applications of Halogen Bonding in Medicinal Chemistry and Chemical Biology. J. Med. Chem. 56, 1363–1388 (2013).

19. Salwiczek, M., Nyakatura, E. K., Gerling, U. I. M., Ye, S. C Koksch, B. Fluorinated amino acids: compatibility with native protein structures and effects on protein–protein interactions. Chem. Soc. Rev. 41, 2135–2171 (2012).

20. Jumper, J. et al. Highly accurate protein structure prediction with AlphaFold. Nature 5G6, 583–589 (2021).

21. Baek, M. et al. Accurate prediction of protein structures and interactions using a three-track neural network. Science 373, 871–876 (2021).

22. Rettie, S. A. et al. Cyclic peptide structure prediction and design using AlphaFold2. Nat. Commun. 16, 4730 (2025).

23. Zhang, C. et al. HighFold: accurately predicting structures of cyclic peptides and complexes with head-to-tail and disulfide bridge constraints. Brief. Bioinform. 25, bbae215 (2024).

24. Anishchenko, I. et al. De novo protein design by deep network hallucination. Nature 600, 547–552 (2021).

25. Lin, H. et al. HighPlay: Cyclic Peptide Sequence Design Based on Reinforcement Learning and Protein Structure Prediction. J. Med. Chem. 68, 12047–12057 (2025).

26. Bryant, P. C Elofsson, A. EvoBind: in silico directed evolution of peptide binders with AlphaFold. 2022.07.23.501214 Preprint at 10.1101/2022.07.23.501214 (2022).

27. Li, Q., Vlachos, E. N. C Bryant, P. Design of linear and cyclic peptide binders from protein sequence information. Commun. Chem. 8, 211 (2025).

28. Wang, F. et al. Reinforcement Learning-Based Target-Specific De Novo Design of Cyclic Peptide Binders. J. Med. Chem. 68, 17287–17302 (2025).

29. Norn, C., et al. Protein sequence design by conformational landscape optimization. Proc. Natl. Acad. Sci. 118, e2017228118 (2021).

30. Pacesa, M. et al. One-shot design of functional protein binders with BindCraft. Nature 646, 483–492 (2025).

31. Kosugi, T. C Ohue, M. Solubility-Aware Protein Binding Peptide Design Using AlphaFold. Biomedicines 10, 1626 (2022).

32. Li, Q., et al. RareFold: Structure prediction and design of proteins with noncanonical amino acids. 2025.05.19.654846 Preprint at 10.1101/2025.05.19.654846 (2025).

33. Zhu, C. et al. Predicting the structures of cyclic peptides containing unnatural amino acids by HighFold2. Brief. Bioinform. 26, bbaf202 (2025).

34. Hu, H. et al. HFGuidedDesign: de novo design of cyclic peptide binders via structure-guided discrete diffusion. Chem. Sci. 17, 15550–15565 (2026).

35. Rettie, S. A. et al. Accurate de novo design of high-affinity protein-binding macrocycles using deep learning. Nat. Chem. Biol. 1–9 (2025) doi:10.1038/s41589-025-01929-w.

36. Powers, A. C., Janthana, Y. C Hosseinzadeh, P. CyclicMPNN: stable cyclic peptide sequence generation. Digit. Discov. https://doi.org/10.1039/d6dd00083e (2026) doi:10.1039/d6dd00083e.

37. Xu, W. et al. HighMPNN: A Graph Neural Network Approach for Structure-Constrained Cyclic Peptide Sequence Design. IEEE J. Biomed. Health Inform. 1–11 (2025) doi:10.1109/JBHI.2025.3620163.

38. Zhang, C. et al. CycleDesigner: Leveraging CycRFdiffusion and HighFold to Design Cyclic Peptide Binders for Specific Targets. J. Chem. Inf. Model. 65, 6155–6165 (2025).

39. Bhardwaj, G. et al. Accurate de novo design of membrane-traversing macrocycles. Cell 185, 3520–3532.e26 (2022).

40. Hosseinzadeh, P. et al. Anchor extension: a structure-guided approach to design cyclic peptides targeting enzyme active sites. Nat. Commun. 12, 3384 (2021).

41. Salveson, P. J. et al. Expansive discovery of chemically diverse structured macrocyclic oligoamides. Science 384, 420–428 (2024).

42. Hosseinzadeh, P. et al. Comprehensive computational design of ordered peptide macrocycles. Science 358, 1461–1466 (2017).

43. Bhardwaj, G. et al. Accurate de novo design of hyperstable constrained peptides. Nature 538, 329–335 (2016).

44. Mulligan, V. K. et al. Computational design of mixed chirality peptide macrocycles with internal symmetry. Protein Sci. 2G, 2433–2445 (2020).

45. Zhu, Q., Mulligan, V. K. C Shasha, D. Heuristic energy-based cyclic peptide design. PLOS Comput. Biol. 21, e1012290 (2025).

46. Abramson, J. et al. Accurate structure prediction of biomolecular interactions with AlphaFold 3. Nature 630, 493–500 (2024).

47. Krishna, R. et al. Generalized biomolecular modeling and design with RoseTTAFold All-Atom. Science 384, eadl2528 (2024).

48. Cao, S. et al. Accurate structure prediction of cyclic peptides containing unnatural amino acids using HighFold3. Brief. Bioinform. 26, bbaf488 (2025).

49. Lin, H. et al. HighPlay2: Structure-guided design of cyclic peptide candidates containing non-canonical amino acids. Eur. J. Med. Chem. 318, 119160 (2026).

50. Xie, X., Li, C. Z., Lee, J. S. C Kim, P. M. CyclicBoltz1, fast and accurately predicting structures of cyclic peptides and complexes containing non-canonical amino acids using AlphaFold 3 Framework. 2025.02.11.637752 Preprint at 10.1101/2025.02.11.637752 (2025).

51. Cho, Y., Pacesa, M., Zhang, Z., Correia, B. E. C Ovchinnikov, S. Boltzdesign1: Inverting All-Atom Structure Prediction Model for Generalized Biomolecular Binder Design. 2025.04.06.647261 Preprint at 10.1101/2025.04.06.647261 (2025).

52. Grambow, C. A., Weir, H., Cunningham, C. N., Biancalani, T. C Chuang, K. V. CREMP: Conformer-rotamer ensembles of macrocyclic peptides for machine learning. Sci. Data 11, 859 (2024).

53. Berman, H. M. et al. The Protein Data Bank. Nucleic Acids Res. 28, 235–242 (2000).

54. Gazizov, A. et al. AF2BIND: predicting small-molecule binding sites using the pair representation of AlphaFold2. Nat. Methods 1–10 (2026) doi:10.1038/s41592-026-03011-2.

55. Koehl, A., Jagota, M., Erdmann-Pham, D. D., Fung, A. C Song, Y. S. Transferability of Geometric Patterns from Protein Self-Interactions to Protein-Ligand Interactions. Pac. Symp. Biocomput. Pac. Symp. Biocomput. 27, 22–33 (2022).

56. Polizzi, N. F. C DeGrado, W. F. A defined structural unit enables de novo design of small-molecule–binding proteins. Science 36G, 1227–1233 (2020).

57. Tsaban, T. et al. Harnessing protein folding neural networks for peptide–protein docking. Nat. Commun. 13, 176 (2022).

58. Ko, J. C Lee, J. Can AlphaFold2 predict protein-peptide complex structures accurately? 2021.07.27.453972 Preprint at 10.1101/2021.07.27.453972 (2021).

59. Chang, L. C Perez, A. Ranking Peptide Binders by Affinity with AlphaFold. Angew. Chem. Int. Ed. 62, e202213362 (2023).

60. Wohlwend, J. et al. Boltz-1 Democratizing Biomolecular Interaction Modeling. 2024.11.19.624167 Preprint at 10.1101/2024.11.19.624167 (2025).

61. Passaro, S. et al. Boltz-2: Towards Accurate and Efficient Binding Affinity Prediction. 2025.06.14.659707 Preprint at 10.1101/2025.06.14.659707 (2025).

62. Lipinski, C. A., Lombardo, F., Dominy, B. W. C Feeney, P. J. Experimental and computational approaches to estimate solubility and permeability in drug discovery and development settings. Adv. Drug Deliv. Rev. 23, 3–25 (1997).

63. Robson-Tull, J. C Rodrigues, J. P. G. L. M. Accurate Physics-Based Flexible Docking of Macrocyclic Ligands. J. Med. Chem. https://doi.org/10.1021/acs.jmedchem.5c03392 (2026) doi:10.1021/acs.jmedchem.5c03392.

64. Mendez, D., et al. ChEMBL: towards direct deposition of bioassay data. Nucleic Acids Res. 47, D930–D940 (2019).

65. Silvestri, A. P. et al. DNA-Encoded Macrocyclic Peptide Libraries Enable the Discovery of a Neutral MDM2–p53 Inhibitor. ACS Med. Chem. Lett. 14, 820–826 (2023).

66. Huang, Y. et al. Discovery of Highly Potent p53-MDM2 Antagonists and Structural Basis for Anti-Acute Myeloid Leukemia Activities. ACS Chem. Biol. G, 802–811 (2014).

67. Roehrig, S. et al. Discovery of the Novel Antithrombotic Agent 5-Chloro-N-({(5S)-2-oxo-3-[4-(3-oxomorpholin-4-yl)phenyl]-1,3-oxazolidin-5-yl}methyl)thiophene-2-carboxamide (BAY 59-7939): An Oral, Direct Factor Xa Inhibitor. J. Med. Chem. 48, 5900– 5908 (2005).

68. Estrada-Ortiz, N. et al. Artificial Macrocycles as Potent p53–MDM2 Inhibitors. ACS Med. Chem. Lett. 8, 1025–1030 (2017).

69. Škrinjar, P., Eberhardt, J., Tauriello, G., Schwede, T. C Durairaj, J. Have protein-ligand cofolding methods moved beyond memorisation? 2025.02.03.636309 Preprint at 10.1101/2025.02.03.636309 (2025).

70. Masters, M. R., Mahmoud, A. H. C Lill, M. A. Investigating whether deep learning models for co-folding learn the physics of protein-ligand interactions. Nat. Commun. 16, 8854 (2025).

71. Leman, J. K. et al. Macromolecular modeling and design in Rosetta: recent methods and frameworks. Nat. Methods 17, 665–680 (2020).

72. O’Boyle, N. M. et al. Open Babel: An open chemical toolbox. J. Cheminformatics 3, 33 (2011).

73. van Kempen, M. et al. Fast and accurate protein structure search with Foldseek. Nat. Biotechnol. 42, 243–246 (2024).

74. Schüttel, M. et al. Solid-phase peptide synthesis in 384-well plates. J. Pept. Sci. 30, e3555 (2024).

75. Nielsen, A. L. et al. Large Libraries of Structurally Diverse Macrocycles Suitable for Membrane Permeation. Angew. Chem. Int. Ed. 63, e202400350 (2024).

76. Perzborn, E., Heitmeier, S., Buetehorn, U. C Laux, V. Direct thrombin inhibitors, but not the direct factor Xa inhibitor rivaroxaban, increase tissue factor-induced hyperco-agulability in vitro and in vivo. J. Thromb. Haemost. 12, 1054–1065 (2014).

## References

1. Robson-Tull, J. & Rodrigues, J. P. G. L. M. Accurate Physics-Based Flexible Docking of Macrocyclic Ligands. J. Med. Chem. https://doi.org/10.1021/acs.jmedchem.5c03392 (2026) doi:10.1021/acs.jmedchem.5c03392.

2. Westbrook, J. D. et al. The chemical component dictionary: complete descriptions of constituent molecules in experimentally determined 3D macromolecules in the Protein Data Bank. Bioinformatics 31, 1274–1278 (2015).

3. Habeshian, S. et al. Synthesis and direct assay of large macrocycle diversities by combinatorial late-stage modification at picomole scale. Nat. Commun. 13, 3823 (2022).

